# Structural Studies of Nedicistrovirus IRES-Driven, Initiation Factor-independent Translation Shed Light on Key Steps of Eukaryotic Translation Elongation

**DOI:** 10.1101/2025.09.29.679340

**Authors:** Swastik De, Clara G. Altomare, Irina S. Abaeva, Prikshat Dadhwal, Priyanka Garg, Francisco Acosta-Reyes, Zuben P. Brown, Tatyana V. Pestova, Christopher U.T. Hellen, Joachim Frank

## Abstract

We utilized the Nedicistrovirus (NediV) intergenic region (IGR) IRES-mediated, initiation factor-independent translation initiation system and determined high-resolution structures of 80S ribosome complexes with the NediV IRES in various functional states, including binary complexes, aminoacyl-tRNA-bound complexes, and complexes with elongation factor eEF2. In binary complexes, the NediV IRES primarily occupies the ribosomal P site, exhibiting conformational flexibility and engaging the ribosome at multiple interaction sites. Upon translocation, the IRES undergoes structural rearrangements, including destabilization of its PKI domain, facilitating the transition to canonical elongation. Crucially, we captured an eEF2-bound complex, along with an eEF1A-bound post-proofreading complex featuring a mismatched tRNA, the latter representing the first instance of a canonical elongation complex visualized in the presence of a natural, hydrolysable nucleotide and without the addition of any trapping agents. These findings provide a comprehensive structural overview of IGR IRES-mediated translation initiation and its transition to elongation, revealing key mechanistic details of viral translation and proofreading.

## INTRODUCTION

Translation is the process through which ribosomes decode the genetic instructions in mRNA to assemble amino acids into polypeptides. Initiation of translation requires the precise positioning of the initiator aminoacyl-tRNA and the start codon of the messenger RNA (mRNA) in the ribosomal P site. In bacteria, this process is relatively simple, requiring just three initiation factors and registration on the ribosome of the mRNA by a short Shine-Dalgarno sequence near its 5′ end. In contrast, eukaryotic translation initiation is far more complex, involving more than a dozen eukaryotic initiation factors (eIFs) to recruit the ribosome and initiate translation from an AUG start codon via a cap-dependent scanning mechanism [1–3]. The eukaryotic mRNA 5′ cap structure, 7-methyl-guanosine-(5’)-ppp-(5’)N, facilitates ribosome recruitment through the eIF4F cap-binding complex, enabling the 40S ribosomal subunit, associated with eIFs 1, 1A, and 3, as well as the ternary complex formed by eIF2, GTP, and Met-tRNA_i_^Met^, to scan for the start codon.

However, many viruses circumvent host translational control by utilizing internal ribosomal entry sites (IRES), structured RNA elements that enable 5′ end-independent translation, particularly under conditions where cap-dependent translation is compromised, such as viral infection or cellular stress [4,5]. Different viral IRESs vary in their dependency on canonical initiation factors. Among them, the dicistronic RNA genomes of dicistroviruses contain an IRES in the intergenic region (IGR) that mediates initiation of translation of the downstream open reading frame (ORF) 2 by an exceptionally streamlined mechanism. IGR IRESs (recently termed type 6 IRESs [6]) bind directly to ribosomal 40S subunits and/or 80S ribosomes, and translation proceeds from a defined location without the involvement of initiator tRNA or eIFs [7,8]. Instead, ribosomes enter directly into the elongation phase of translation [9, 10]. Translation mediated by IGR IRESs is therefore resistant to, or even stimulated by, regulatory mechanisms and pharmacological compounds that inhibit the canonical initiation process [7, 11,12]. The ability of IGR IRESs to bypass the conventional initiation process renders them a useful tool both for the synthesis of biologics in eukaryotic cell-free translation systems (e.g. [13–17]) and to facilitate characterization of the mechanism and inhibitors of downstream steps in eukaryotic translation, including elongation [9,18,19] and termination [20–22].

Type 6 IRESs form several subgroups, each with shared structural elements and sequence motifs that underlie differences in mechanism of action. Type 6a IGR IRESs, the first to be characterized, are exemplified by Cricket paralysis virus (CrPV). They are ∼190 nt long and consist of three pseudoknot (PK) domains. The nested domain 1 (PKII) and domain 2 (PKIII) are principally responsible for the affinity of ribosomal binding: the conserved internal L1.1a and L1.1b loops in domain 1 bind to the L1 stalk of the ribosomal 60S subunit, and the SL2.1 and SL2.3 stem-loops in domain 2 bind to ribosomal proteins eS25 and uS7, close to the ribosomal E site and the mRNA exit channel [23,24]. Domain 3 (PKI) binds in the decoding center of the 40S ribosomal A site, placing the adjacent region of ORF2 into the mRNA-binding channel [24]. PKI consists of an element that mimics a tRNA anticodon stem loop (ASL) engaged in codon-anticodon interaction with an mRNA triplet, and a conformationally dynamic loop (L3.2) known as the variable region loop (VRL) [25] that influences subsequent translocation steps [26]. In a process termed pseudo-translocation, eEF2 brings PKI from the ribosomal A to the P site and the first codon of ORF2 into the A site; however, this state is meta-stable and the ribosome can undergo spontaneous back-translocation if the codon in the A site is not decoded [9,20,27,28]. Pseudo-translocation is followed by delivery of a cognate aminoacyl-tRNA to the ribosome in complex with eEF1A and GTP (the elongation factor 1·GTP·aa-tRNA ‘ternary complex’ (TC)), release of the aminoacyl-tRNA into the A site, its accommodation, and its eEF2-mediated translocation to the ribosomal P site. This final stage is accompanied by partial unwinding of PKI, such that the ASL-like element relocates to a position equivalent to that of the acceptor stem of E-site tRNA [29].

There are several other distinct structural subclasses of type 6 IRES, the best-characterized of which (type 6b, exemplified by Taura syndrome virus (TSV) and Israeli acute paralysis virus (IAPV)) contain divergent motifs in L1.1a and L1.1b loops and an additional, functionally important stem-loop in domain 3 [30,31]. Type 6b IRESs are thought to promote adoption by the 40S-IRES complex of a conformation that facilitates ribosomal subunit joining [32] and to influence reading frame selection [33]. Specific motifs and structures therefore determine specific aspects of function, such as attachment to the ribosomal A site, mimicry of the ASL/codon interaction, engagement with the L1 stalk, and facilitation of subunit joining. Recently identified subclasses of type 6 IRESs lack one or more structural elements that are typical of type 6a and type 6b IRESs and utilize even simpler mechanisms for initiation [34–36]. For example, type 6d IRESs, epitomized by Nedicistrovirus (NediV), are only ∼165 nt long, and while they also consist of three pseudoknot domains, they lack a SL2.3 stem-loop and canonical motifs in the L1.1a/L1.1b loops [35]. They bypass the entire canonical initiation process, like type 6a and 6b IRESs, but also bypass the subsequent pseudo-translocation step, and instead bind directly to the P site, placing the first codon of the downstream ORF in the A site, where it is accessible for decoding by the TC. Once initiation is complete, the structured IRES domains dissociate from the ribosome, leaving only the ORF mRNA for the ribosome to transition into the canonical elongation cycle.

In each elongation cycle, eEF1A within the TC delivers the incoming aa-tRNA to the A site of the ribosome. Non-cognate codon-anticodon mismatches result in the immediate rejection of the TC, while near-cognate and cognate pairings trigger GTP hydrolysis on eEF1A, leading to its release and the accommodation of the aa-tRNA into the A site [37]. Following this step, eEF2, bound to GTP, triggers intersubunit rotation, a rotation of the small subunit relative to the large subunit [38, 39], and thereby translocates the tRNAs together with their bound mRNA (“tRNA2-mRNA”) from the P and A sites into the hybrid P/E and A/P sites, and eventually into the E and P sites. The spatial precision of this process is essential for maintaining high translational fidelity.

Although spontaneous intersubunit rotation does occur, leading to translocation of the tRNA2-mRNA moiety, eEF2·GTP accelerates this process [19]. By binding to the A site, eEF2 induces specific ribosomal conformational changes, thereby ensuring maintenance of the reading frame and enforcing translational fidelity [40,41]. The ribosome is prone to frameshifting during this phase, but the diphthamide modification of eEF2 helps prevent these errors by stabilizing the codon in its correct position [38,40,42].

Published structures of eEF2-bound ribosomes span a range of species, from yeast (*Saccharomyces cerevisiae*) (e.g., 7OSM) to mammals like rabbit (*Oryctolagus cuniculus*, PDB IDs 6MTD and 6MTE) and humans (PDB IDs 6Z6M), with resolutions ranging from 11.7Å to 3.0Å [42,43,44]. These structures capture ribosomes in different eEF2-bound conformations, including rotated and non-rotated states (such as in PDB IDs 6MTE and 6MTD, bound to SERBP1 and GDP) as well as those with ligands like GDPNP (PDB ID 2P8W) and GDPCP (PDB ID 7OSM) [38,42]. Additional structures of ribosome-eEF2 complexes include Nelfinavir-treated neuronal ribosomes (PDB ID 7LS2), Puromycin-treated ribosomes with EBP1, SERBP1, and eEF2 (PDB ID 6Z6M), and viral IRES-bound ribosomes (PDB ID 5IT7 and 6D9J) [29,44,45,46]. Crucially, only one of these structures of eEF2-bound ribosomes [29] formed under natural reaction conditions, and all other structures correspond to complexes assembled in the presence of non-native nucleotides, antibiotics, or biochemical inhibitors.

Only a limited number of published structures have captured complexes with eEF1A, including those stabilized by translation inhibitors such as Didemnin B (PDB ID 5LZS) or Angiogenin (PDB ID 9BDP) [47,48], as well as one obtained through in situ imaging of yeast lamellae (PDB ID 8Z71) [49].

Current knowledge of elongation cycle dynamics relies heavily on these static structures, which represent ribosome complexes stabilized in specific states using GTP analogs or other chemical means. However, they leave considerable gaps in knowledge as they do not include short-lived intermediate states instrumental for ensuring fidelity in eukaryotic translation. This study used cryo-EM to determine the structures of rabbit ribosomal initiation and early elongation complexes assembled on the NediV IRES. Included is the visualization of one previously unreported intermediate state in the mammalian translocation cycle: the eEF1A-bound ribosome with A*/T-tRNA stalled at a codon-anticodon mismatch. Its structure reveals a distortion in the tRNA anticodon loop and unusual eEF1A positioning, as well as offering insights into the structure of the TC in this state. Notably, this study characterizes biologically relevant complexes on pathway bearing GTP, validating previous conclusions drawn from kinetically trapped complexes.

The structures we present form a detailed molecular timeline of IGR IRES-mediated translation initiation and its seamless handoff to canonical elongation. It is seen in intriguing detail how the NediV IRES directly binds to the 80S ribosome’s P site, bypassing pseudo-translocation, and thereby establishes a stable platform for accurate initiation. Our structural snapshots illuminate how IRES dynamics go hand in hand with ribosomal conformational changes and the binding of elongation factors in enabling efficient translation. Most importantly, we observe a novel post-proofreading eEF1A-bound complex, captured without chemical trapping, that highlights a mechanistic basis for fidelity control. These insights not only clarify the NediV IRES’s unique strategy but also shed light on the nature of conserved translational checkpoints, underscoring the utility of IGR IRESs in dissecting translation mechanisms.

## METHODS

### Plasmids

The transcription vector for rabbit cytoplasmic tRNA^Ala^ (GenBank accession no. Y17041), and expression vectors for eRF1 and a truncated form of eRF3 lacking a.a. 1–138 have been described [9,50]. Transcription vectors for NediV IGR IRES-containing mRNAs were made by GeneWiz (South Plainfield, NJ). pUC57-NediV (wt) [35]contains a stable hairpin (5′-GGCTCGAGGCCCGGTGACGGGCCTCGGGCC-3′ [ΔG=−32.70 kcal/mol]) and NediV nt 1161–1606, inserted downstream from a T7 promoter between BamH1 and XbaI sites of pUC57. The second codon (ACA (Thr)) of the open reading frame was substituted by a UAA stop codon in the NediV-[Ala.STOP] variant. (Figure S3D)

### Purification of factors, ribosomal subunits, and aminoacylation of tRNA

Native eEF1H, eEF2, total aa-tRNA synthetases and 40S and 60S ribosomal subunits were purified from rabbit reticulocyte lysate (Green Hectares, Oregon, WI) [51,52]. eRF1 and eRF3 were purified after expression in *Escherichia coli* [50]. mRNAs and tRNA were transcribed using T7 RNA polymerase (Thermo Fisher). In vitro transcribed tRNA^Ala^ was refolded and then aminoacylated using total native aminoacyl-tRNA synthetases [9].

### Assembly and analysis of ribosomal complexes

Binary IRES/80S ribosome complexes were assembled by incubating 7 pmol NediV (wt) mRNA for 5 min at 37°C in 40ul reaction volumes that contained buffer A [2 mM dithiothreitol (DTT), 100 mM potassium chloride, 20 mM Tris (pH 7.5), 2.5 mM magnesium chloride and 0.25 mM spermidine] supplemented with 3.2 pmol 40S subunits and 5 pmol 60S subunits. Ribosomal elongation complexes were assembled by incubating 7 pmol NediV (wt) or NediV-[Ala.STOP] mRNA for 5 min at 37°C in 40ul reaction volumes that contained buffer A supplemented with 1 mM GTP and 3.2 pmol 40S subunits, 5 pmol 60S subunits, 7pmol Ala-tRNA^Ala^, 10 pmol eEF1H, 15 pmol eEF2, 15 pmol eRF1 and 20 pmol eRF3. Assembly and translocation of complexes was confirmed by toe-printing using avian myeloblastosis virus (AMV) reverse transcriptase (Promega) and a cognate primer as described [35].

### Sample Preparation and Data Collection

For (A) the NediV-IRES:80S complex (binary setup), (B) the NediV-IRES·80S·tRNA·eEF2 complex (elongation setup) and (C) the NediV-IRES·80S·tRNA·eRF1 complex (termination setup), 3 μl of ribosomal complexes at concentrations of (A) 500 nM and (B, C) 80 nM were applied to plasma-treated, thin carbon-coated holey carbon grids (QUANTIFOIL R1.2/1.3) [53], blotted for (A) 5 seconds and (B, C) 9 seconds, and rapidly frozen in liquid ethane using an FEI Vitrobot. Following freezing, the grids were transferred to a Polara-G2 microscope, operating at 300 kV and equipped with a Gatan K2 Summit direct detector. For the NediV-IRES·80S complex, 10,690 frame stacks were captured in counting mode with a dose rate of 8e^−^/pix/s at a magnification of 31,000x, corresponding to a calibrated pixel size of 1.24 Å. For the NediV-IRES·80S·tRNA·eEF2 complex, 4,193 frame stacks were recorded in counting mode with a dose rate of 8e^−^/pix/s at a magnification of 23,000x, yielding a calibrated pixel size of 1.66 Å.

For the NediV-IRES·80S·tRNA·eRF1 complex, 3,273 frame stacks were collected in counting mode at a dose rate of 16e^−^/pix/s at a magnification of 39,000x, with a calibrated pixel size of 0.95 Å. Frame stacks for all complexes were recorded in automatic mode using Leginon software [54], which determined that defocus values ranged (A) from 2.5 to 3.5 μm and (B, C) from 1.5 to 3.5 μm. Frame alignment was performed using Motioncor2 [55], and data collection was monitored in real-time using APPION [56].

### Data Processing and Classification

For all ribosomal complexes, aligned micrographs were manually screened for quality. Contrast transfer function (CTF) parameters were estimated using GCTF [57]. Particle picking was performed using Gaussian picking in GAUTOMATCH, with a particle diameter set to 350 Å. Subsequent 2D and 3D classifications and refinements were carried out using RELION [58,59]. The picked particles were binned four times and subjected to 2D classification to separate 80S ribosomes from non-ribosomal particles and noise. Initial consensus models for the 80S particle sets were generated using the 3D Refine procedure.

To further clean high-noise datasets, a global 3D classification was performed. A mask enclosing the ∼1/3 of the 40S, IRES, and 60S was created to prioritize IRES density and applied in RELION for separate 3D classification of each dataset. The resulting classes were divided into IRES-containing complexes and E-site tRNA-containing complexes. The latter, as they are no longer bound to the IRES but retain the linked ORF portion of the viral mRNA, can be considered elongation complexes on the canonical pathway. High-resolution reconstructions of IRES-containing complexes without tRNA or additional factors were also obtained. However, complexes containing tRNA and eEF2 exhibited noisy densities and image defects, defying further 3D classification.

Masked refinements were performed on the unbinned data for selected classes using RELION, followed by non-uniform refinement in cryoSPARC [60] to improve resolution in flexible regions. Despite these refinements, factor densities remained poorly resolved. To better address dataset heterogeneity, cryoSPARC variability analysis and CryoDRGN [61] were employed.

Following two rounds of particle filtering, CryoDRGN heterogeneity analysis of the NediV-IRES:80S dataset yielded clean volumes, primarily consisting of 80S ribosomes with P-site-loaded IRES, 80S ribosomes with A-site-loaded IRES, and empty 80S ribosomes. For the NediV-IRES·80S·tRNA·eEF2 dataset, CryoDRGN heterogeneity analysis identified high-resolution classes, including 80S with a non-translocated IRES and A-site tRNA, 80S with a singly translocated IRES and P-site tRNA, 80S with a singly translocated IRES, P-site tRNA, and eEF2, and 80S in a canonical elongation complex with eEF1A and A/T* tRNA. Finally, the NediV-IRES·80S:tRNA·eRF1 dataset yielded multiple termination complexes (to be presented elsewhere) as well as a canonical elongation complex containing eEF2 and P-site tRNA.

### Model Building

Atomic models were built for the major complexes:

A. the NediV-IRES·80S binary complexes,
B. the NediV-IRES·80S·tRNA complex,
C. the NediV-IRES·80S·tRNA·eEF2 complexes, and
D. the NediV-IRES·80S·tRNA·eEF1A complex.

For the binary complex (A), the 80S ribosome from PDB 5LZS was used as the initial model. For (B), (C), and (D), the starting models were 5LZS (ribosome and eEF1A) and 6GZ5 (ribosome, eEF2 and tRNAs) respectively. The rRNA sequences were changed for rabbit rRNA based on the study by De et al., 2025 [62]. The NediV IRES was modeled de novo in Coot [63], guided by its predicted secondary structure [35]. All atomic models were further adjusted manually in Coot and ChimeraX [64] to optimize fit into the experimental density maps. Final refinements were performed using Phenix real-space refine [65], with appropriate secondary structure restraints and validation metrics monitored throughout.

## RESULTS

### Cryo-EM analysis of ribosomes bound to the NediV IRES

Cryo-EM analysis of binary NediV IRES·80S ribosome complexes revealed two distinct classes displaying a canonical non-rotated ribosomal state (i.e., without intersubunit rotation) with domain 3 (PKI) bound to either the ribosomal P site or to the A site (Figure 1A, 1B, S1, S2A, S2B). The resolution of these classes was 4.2 and 4.8 Å, respectively (Supp. Table 1). As with other type 6 IRESs, NediV IRES·80S complexes are conformationally dynamic and sample different rotational states [34,66,67,68]. Another IRES-containing class with a rotated 60S subunit was obtained, but the IRES density was too noisy for model building or further analysis (Figure S3A-C).

**Figure 1.**
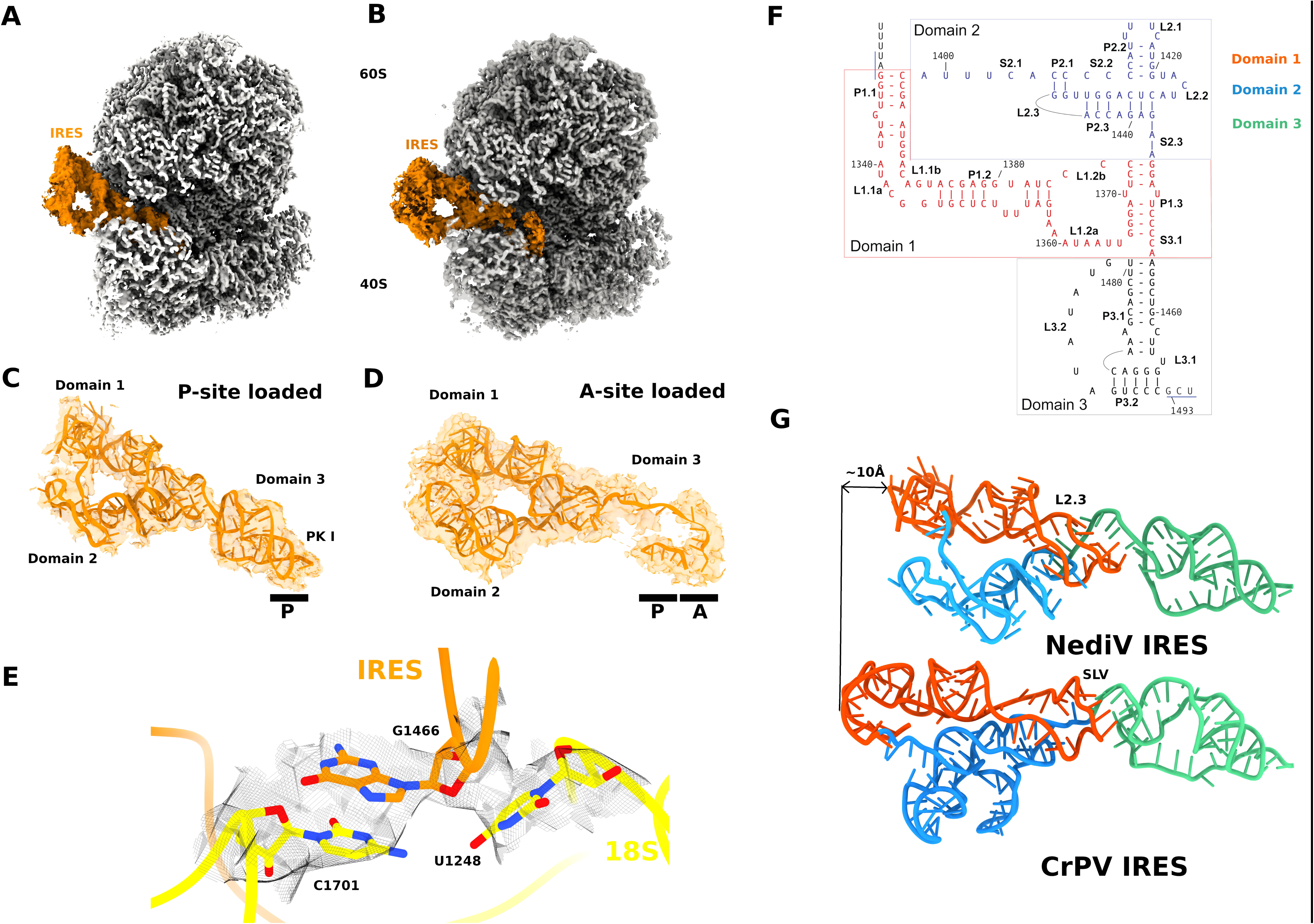
Structural Features of the NediV IRES in the Initiation Complex. (A) and (B) Cryo-electron microscopy (cryo-EM) density maps of the NediV IRES (dark orange) bound to the mammalian 80S ribosome (grey). The maps reveal two distinct conformational classes: the more abundant P-site-loaded state (A) and the less-stable A-site-loaded state (B). (C) and (D) Zoomed-in views of the NediV IRES within the ribosomal mRNA cleft, showing its distinct binding positions. Panel (C) highlights the well-ordered IRES occupying the P site in a canonical, non-rotated ribosomal state. Panel (D) shows the less-ordered IRES in the A site, which appears to block access to both the A and P sites. (E) Zoomed in view of the P-site-bound IRES and 18S rRNA residues at the A-gate, demonstrating the high-quality resolution of the map in this region, which enabled detailed structural analysis of the IRES-ribosome interactions. (F) Predicted secondary structure of the full-length NediV IRES, color-coded to delineate its characteristic domains, including the nested pseudoknots (PKII and PKIII) and the independently folded PKI. (G) A structural comparison of the NediV IRES (type 6d) and CrPV IRES (type 6a) (PDB ID 6D90) showing the length difference between the IRESs when the PKI for both the IRESs were aligned. Same color scheme as (F) was used.

The ribosome-bound NediV IRES has the characteristic architecture of type 6 IRESs, comprising the nested PKII and PKIII and the independently folded PKI (Figure 1F; Figure S1). The NediV PKI domain closely resembles PKI of the type 6a CrPV IRES, which comprises helix P3.1 (11 base-pairs (bp) and one unpaired nucleotide), helix P3.2 (5 bp), the 9 nt-long variable region loop (VRL) that connects them, and one unpaired nucleotide. The equivalent elements in the NediV IRES are helix P3.1 (comprising 9 bp and two unpaired nucleotides), helix P3.2 (5 bp), the 7nt-long VRL and one unpaired nucleotide [35]. The principal differences between type 6d NediV-like IRESs and type 6a IRESs are that domain 1 in the former has conserved motifs in loops L1.1a and L1.1b that are unlike those in type 6a IRESs, and domain 2 has a simple loop in place of the SL2.3 stem-loop (Figure 1G). These differences likely account for the different positions of type 6a and 6d IRESs in binary ribosomal complexes, as described below.

Type 6a IRESs bind in the A site [24], whereas PKI is in the P site in the overwhelming majority (∼70%) of binary NediV IRES:80S complexes (Figure 1A, 1C). This location is consistent with toe-printing and other biochemical data, which showed that in elongation-competent complexes, the NediV IRES occupies the P site and the first codon of ORF2 is placed in the A site [35]. Binding of the NediV IRES to the P site is stabilized by three, and possibly four sets of interactions with the ribosome (Figure S2). First, the L1.1 region in domain 1 binds to ribosomal protein uL1 of the 60S subunit and to nucleotides of the L1 stalk (Figure S2B, S2C, S2D). These interactions anchor domain 2 securely to facilitate the insertion of the remainder of the IRES into the mRNA cleft and intersubunit space. PKI binds to the ribosome via residue G1466, which interacts with nt. 1701 of the 18S rRNA at a distance of 7.7 Å, while G1467 is positioned near 18S rRNA nt.1248, with a distance of 5.8 Å (Figure 1E, S4A). G1468 interacts with Arg140 of uS19, maintaining a distance of 9.0 Å, and IRES residues A1498 and A1500 are in close proximity to Gln113 and Phe124 of ribosomal protein uS5 and A1485 to Arg66 of uS11, creating stabilizing interactions on both sides of PKI and place PKI firmly into the ribosomal P site (Figure S4B, S4C). The third set of interactions involves the VRL, which interacts with ribosomal protein uS7 (Figure S2B, S2C). Finally, C1423, in the highly conserved L2.2 loop in domain 2, is in close proximity to eS25 at the head of the 40S subunit (Figure S2E), but details of this interaction could not be established because of the lower resolution of the map in this area. The inability of the NediV IRES to establish a stable interaction with eS25 may be a contributing factor to the weak binding of this IRES to the 40S subunit alone, and to its preferred binding to P site in 80S ribosomes.

The functional importance of each of these binding sites has been confirmed by mutational analysis of the NediV IRES and of the related type 6d Antarctic picorna-like virus (APLV) IRES [35]. Notably, identical or similar interactions have been observed in the binary 80S·HalV IRES complex [34], and in the post-translocation type 6a (CrPV) and type 6b (TSV and IAPV) IRESs [20,32,69]. The NediV IRES forms a complex, by binding directly to the 80S ribosome, that is directly analogous to complexes formed on type 6a and 6b IRESs after they have bound to 80S ribosomes and undergone pseudo-translocation.

A substantially smaller proportion of binary complexes contain the NediV IRES bound to the A site: domain 3 (PKI) in these complexes appears disordered, allowing only a single strand of the IRES entering the IRES to be modeled (Figure 1D). The P site-loaded state thus appears to be more stable than the A site-loaded state, and is well-resolved, ordered and (as described above) forms multiple contacts with the ribosome. Whereas the A site is accessible for binding of aminoacyl-tRNA in the P site-bound state, the NediV IRES binds to the A site of nonrotated ribosomes in a perpendicular position, extending through the P site and effectively blocking access of aminoacylated tRNA to both A and P sites. It is possible that this A site-loaded configuration represents an unstable state that can transition toward the more stable P site-loaded state.

### The structure of 80S**·**NediV IRES**·**aa-tRNA complexes

Decoding of the A-site codon in binary 80S·NediV IRES complexes yields a transient ternary [80S·IRES·aa-tRNA] complex, and although the latter corresponds to a key stage in translation mediated by type 6 IRESs, its structure has not previously been determined. To capture this complex, the NediV wt IRES was incubated with Ala-tRNA^Ala^, eEF1H (a complex of eIF1A and its guanine-nucleotide-exchange factor eEF1B), eEF2, GTP, and mammalian 80S ribosomes pre-assembled from purified 40S and 60S subunits (also defined as ‘elongation setup’ in Methods). This preparation yielded high-quality cryo-EM data, with 4,139 micrographs collected at a resolution of 1.66 Å per pixel. After particle selection and 2D classification, 19,647 particles of 80S ribosomes bound to the IRES and Ala-tRNA^Ala^, but lacking elongation factors or additional tRNAs, were refined into three ribosomal classes, of which only one resolved the tRNA density sufficiently for model building (6.4 Å) (Supp. Table 1). Binding of Ala-tRNA^Ala^ to these complexes was mediated by eEF1H·GTP, and both the absence of this factor from these complexes and the observed full accommodation of Ala-tRNA^Ala^ in the A site (Figure 2A, 2D) are together indicative of direct recognition of the A-site codon by cognate tRNA and consequent triggering of GTP hydrolysis, leading to release of Ala-tRNA^Ala^ and dissociation of eEF1H·GDP.

**Figure 2.**
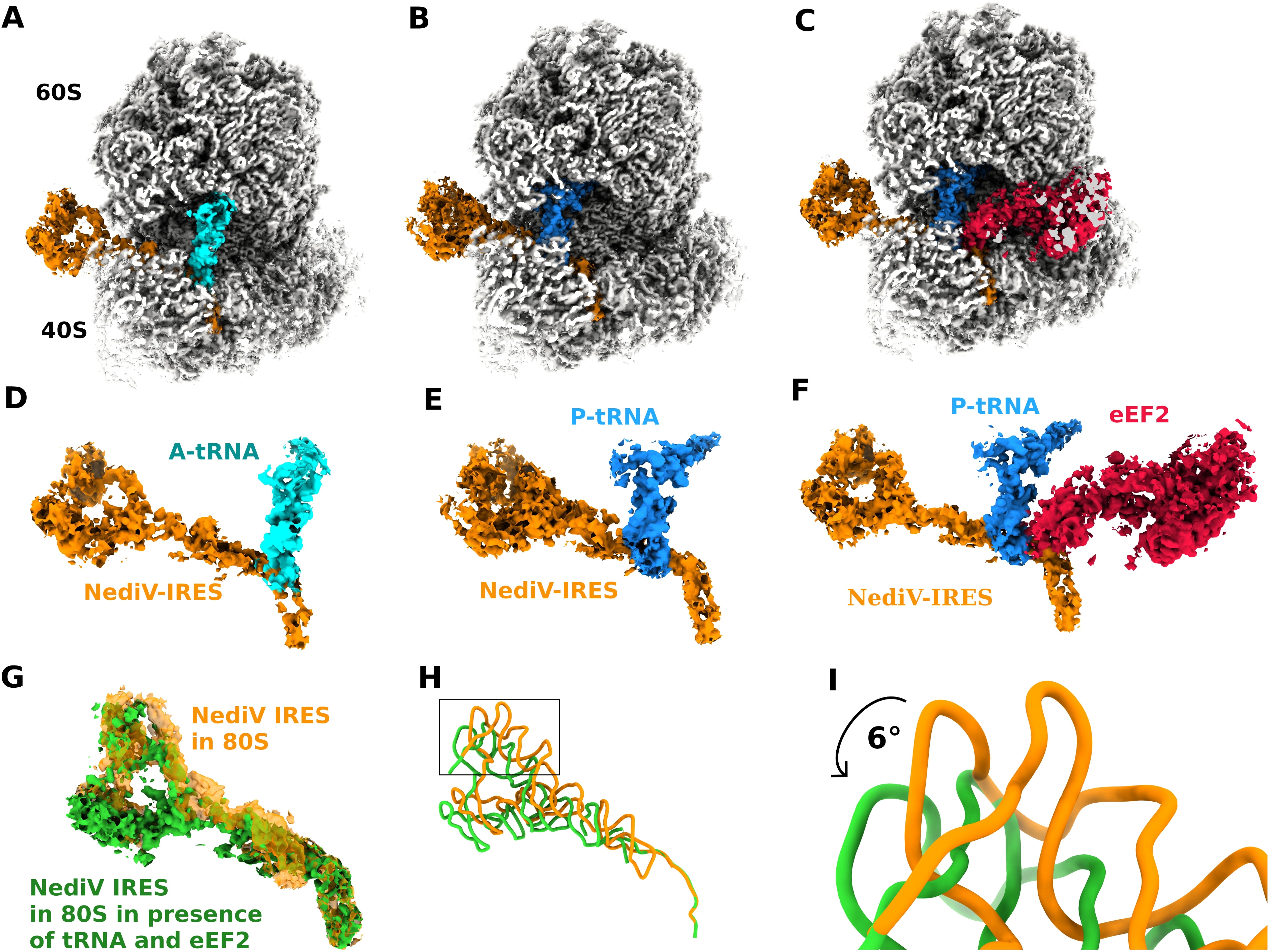
Translation Elongation Complexes on the NediV IRES. (A), (B), and (C) Cryo-EM reconstructions illustrating the transition from initiation to the first elongation step. The NediV IRES is shown in dark orange, with the ribosome in grey. Panel (A) shows the initial ternary complex with Ala-tRNA (cyan) bound in the A site. Panel (B) shows a state where the IRES remains stably bound in the P site after the tRNA has been translocated to the P site. Panel (C) shows the post-translocation complex, featuring eEF2 (crimson) and a translocated P-site tRNA (dodger blue). (D), (E), and (F) Focused views of the complexes shown in (A), (B), and (C), respectively, providing a detailed look at the positioning of the IRES and its interactions with the ribosome, tRNA, and eEF2. (G) and (H) Structural alignment of the binary (80S+IRES) and quaternary (80S+IRES+tRNA+eEF2) complexes, showing the conformational changes of the IRES itself. Panels (G) and (H) illustrate how Domains 2 and 3 of the IRES rotate by approximately 6 degrees relative to their position in the binary complex as the ribosome transitions to the elongation state. (I) A close-up view of IRES Domain 1 models, showing the subtle but critical conformational rearrangements that occur to accommodate the full elongation machinery.

The three structural classes correspond to ribosomal states with varying degrees of slight intersubunit rotation: 1.1°, 1.4° and 1.6°. Despite these rotational differences, the NediV IRES remains stably bound to the ribosomal P site (Figure 2B), showing minimal movement relative to the 18S rRNA. C1243 in the L2.2 loop remains in close proximity to eS25 as in the binary complex (Figure S2E), and there is no significant compression or expansion of domains 1 and 2 in this complex relative to the binary complex (Figure 2D, 2E, 2F). Indeed, the initial binding of Ala-tRNA^Ala^ to the ribosomal A site appears to stabilize the overall ribosome-IRES interaction, particularly enhancing the rigidity of domains 1 and 2 (Figure 2D). These and other interactions observed in the binary complex anchor domains 1 and 2 of the IRES, allowing the ribosome to adopt both canonical and hybrid states, which are required for elongation. This dynamic behavior is consistent with that of type 6a IRES elements, such as CrPV [66], but differs significantly from IAPV and TSV type 6b IRESs, which stabilize the ribosome in the non-rotated state [32,69].

The least-rotated (1.1°) 80S·NediV IRES complex exhibits the best-resolved A-site tRNA; the 1.4°-rotated complex shows the A-site tRNA less well-resolved; and in the 1.6°-rotated complex, the A-site tRNA is poorly resolved even though the IRES is well-defined. The IRES in the Ala-tRNA^Ala^-bound complex is rotated 2–3 degrees away from the L1 stalk relative to its position in the binary 80S·NediV IRES complex. The dynamic behavior of the 40S subunit in these complexes mirrors the dynamics observed in canonical elongation, where tRNA delivery to the ribosomal A site induces conformational changes in the ribosome, including reorientation of G626 and flipping-out of the ‘monitoring’ nucleotides A1824 and A1825 from helix 44 (G530, A1492 and A1493 in *E. coli*), as well as domain closure [47,70,71,72] (Figure S5).

### The structure of the post-translocation complex with eEF2

The 80S·IRES·aa-tRNA complex resembles neither the rotated-1 state [70] nor the rotated-2-Pre state [73], with a meager 1.1-degree rotation observed in our structures. The slightly rotated state observed before forms during the canonical elongation process, except that the P site in the 80S·IRES·aa-tRNA complex is occupied by PKI rather than by aminoacylated tRNA. This complex is thus primed for the binding of eEF2 and subsequent translocation.

To investigate the binding of eEF2 and the ensuing translocation step, we assembled the 80S·IRES·aa-tRNA·eEF2 complex by incubating the NediV IRES with Ala-tRNA^Ala^, eEF1H, eEF2, GTP, and mammalian 80S ribosomes pre-assembled from purified 40S and 60S subunits (defined as the elongation setup in Methods). A total of 4,139 micrographs were collected at 1.66Å/pixel resolution. Heterogenous classification identified one class with 18,360 particles (Supp. Table 1), capturing the ribosome in the non-rotated state. In this state, eEF2 occupies the A site, tRNA has been translocated to the P site, and the IRES has also been singly translocated (Figure 2C, 2F). The position of the 40S subunit head remains largely unchanged in this eEF2/P-site tRNA state compared to the 80S·NediV IRES complex with aa-tRNA in the A site. Detailed structural comparisons show that these complexes closely resemble the recently described POST-3 state of the elongation complex (PDB ID 6GZ5; [40]). The present eEF2-bound structure maintains a near-zero-degree rotation for both the head and body, suggesting that it corresponds to a complex at the end of the translocation cycle, immediately prior to the release of eEF2.

Two key structural differences are apparent between the IRES in this and the preceding two complexes. Domains 2 and 3 have rotated by ∼6 degrees relative to their position in the binary complex, moving close to the 40S head (Figure 2G – 2I). Whereas the P site-bound PKI was stabilized in the presence of A site tRNA (Figure 2D), the translocated PKI in complexes containing P-site tRNA was destabilized, and is visible only up to A1473, 14.8Å from C40 of the tRNA (Figure 3B). The destabilization of NediV PKI in the E site, which is reminiscent of the disassembly of the analogous domain in the CrPV IRES upon translocation from P to E sites [29], may support the threading of the viral ORF2 coding region into the mRNA channel, preparing the ribosome for elongation. The IRES mRNA is well resolved all the way along its path through the ribosome. Past the A site, it follows the mRNA cleft while interacting closely with the 18S rRNA (sandwiched between the residues 1825 and 1701) and ribosomal proteins uS3, and uS5 (Figure S2B, S6A, S6B).

**Figure 3.**
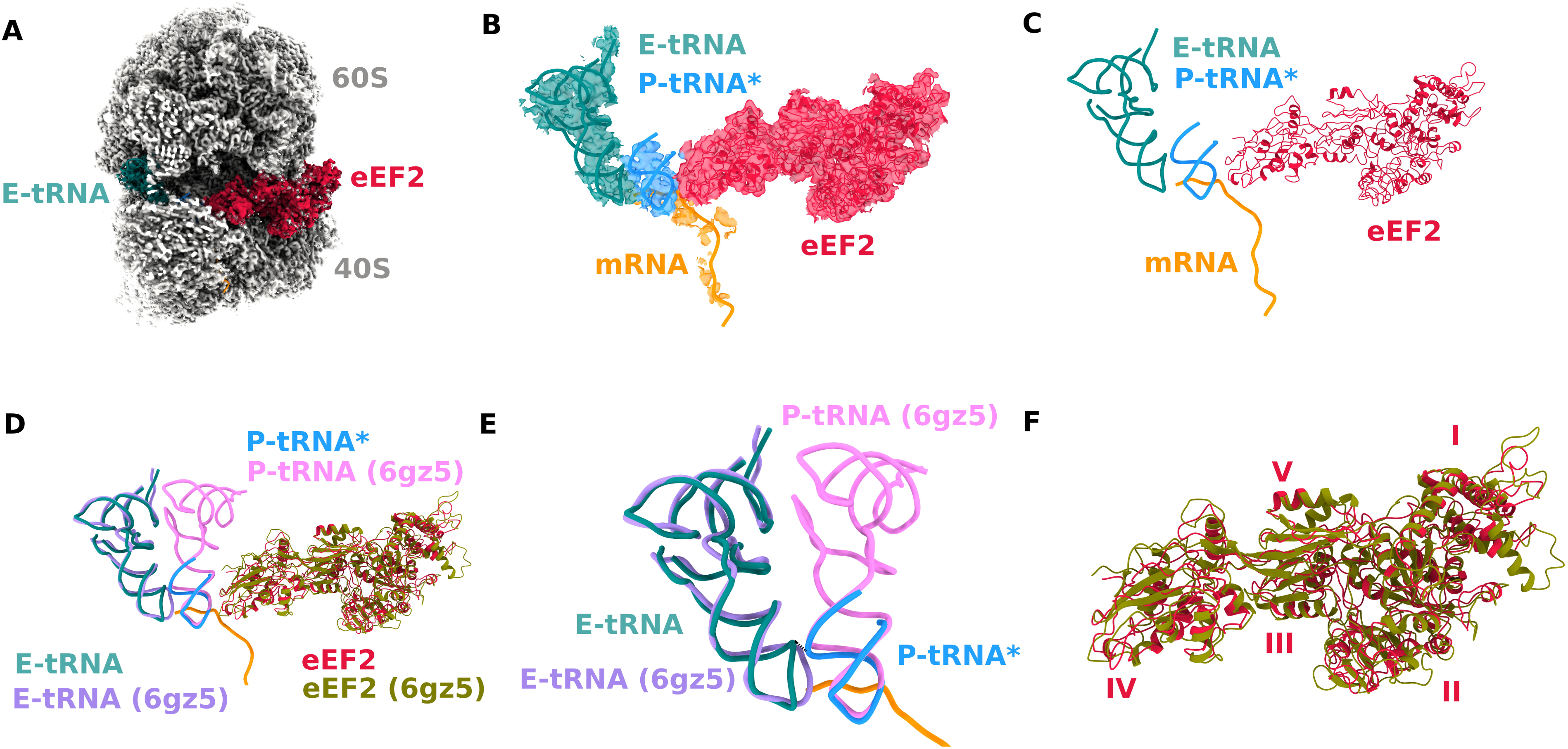
Elongation Complex in the Presence of eEF2. (A) Cryo-EM structure of a fully elongation complex, captured from the termination setup. This state contains a translocated P-site tRNA (dodger blue), an E-site tRNA (teal), and eEF2 (crimson). This class represents the transition from IRES-mediated initiation to canonical elongation. (B) and (C) Focused views of the complex in (A). Panel (B) shows the cryo-EM map, and (C) shows the atomic model, highlighting the arrangement of eEF2 and the two tRNAs. Notably, the P-site tRNA is well-resolved only at its lower half, while the off-pathway E-site tRNA is exceptionally stable, reflecting the strength of the stable occupation of the E site. (D) and (E) Comparative analysis of the tRNAs in the E and P sites. These panels highlight the unusually close proximity between the P-site and E-site tRNAs, with their backbones nearing a steric clash, a novel observation in eukaryotic elongation complexes. (F) A comparative analysis of the eEF2 conformation in our canonical complex versus the POST-3 state (PDB ID 6GZ5).

We also isolated a distinct group of particles from our termination setup experiment (see Methods) in which we could identify eEF2 on the ribosome (Figure 3A). In this class we captured an eEF2-containing 80S ribosome with two tRNAs present and only the ORF mRNA visible (Figure 3B, 3C), marking the transition into a canonical elongation state. This class displays a unique arrangement of eEF2 and the P- and E-tRNAs, with a slight (1.4°) intersubunit rotation and the tRNAs adopting hybrid states, engaging in previously characterized tRNA-mRNA and tRNA-ribosomal interactions [41] (Figure 3A). The lower half of the P-site tRNA is significantly better resolved than the rest of the molecule (Figure 3B, 3C, 3D, 3E), reflecting the strength of the codon–anticodon interaction established after the first translocation step. Although the presence of deacylated tRNA in the E site may reflect binding due to its presence in molar excess over ribosomes during sample preparation (e.g. [70]), stable occupation of the E site by deacylated tRNA is a characteristic of eukaryotic elongation complexes [73,74].

The structural map strongly resembles the classical POST-3 state of elongation, with tRNAs in the P and E sites and the L1 stalk rotated into a stabilized conformation, consistent with observations in bacterial and rabbit elongation complexes [40,41,75]. The subtle tilt of the 40S subunit head (1.4°) could either be a remnant of the prior translocation event or an inherent feature of higher mammalian ribosomes at rest. The conformational adjustments of the 40S subunit are linked to the P-gate dynamics. In our structure, the P gate remains intact, with a gate distance of ∼17Å, suggesting that it halted the transition of tRNAs from hybrid to fully accommodated P/P and E/E positions (Figure 3C).

A particularly striking feature of this class is the unprecedentedly close proximity of the P-site and E-site tRNAs. Across all classes, the E-site tRNA displays exceptional stability and high occupancy, potentially having entered the ribosome via the open L1 stalk. Meanwhile, the P-site tRNA part of the map exhibits lower resolution, with the anticodon arm, T-arm, D-arm, and acceptor stem largely unresolved, indicating weaker stabilization. Notably, the P-site tRNA appears to invade the E site, bringing its backbone residue 40 into close contact with the backbone of the nearest E-site tRNA residue (C36) at an unusually short distance of ∼3 Å, nearing steric clash (Figure S7). Additionally, O2 of U32 in the P-site tRNA is in close proximity with O2 of G35 in the E-site tRNA (Figure 3E and S7).

Both structural classes exhibit nearly identical positioning of eEF2, which interacts with the ribosomal GTPase-associated center (GAC) of the large ribosomal subunit. The ribosome-eEF2 complex undergoes significant conformational rearrangements upon GTP hydrolysis, as observed in 80S·ADPR-eEF2·GDPNP and 80S·ADPR-eEF2·GDP·sordarin structures [38,41]. These rearrangements shift domains G′, I, and II of eEF2 toward the GAC while displacing domains III, IV, and V further away. The Switch 1 loop, a key regulatory element, adopts an ordered conformation in the GTP-bound state, similar to EF-G [76,77], engaging the terminal phosphate groups of GTP via Mg²⁺ coordination. Following GTP hydrolysis, the Switch 1 loop becomes disordered, as confirmed by 80S·ADPR-eEF2·GDP·sordarin structures (PDB ID 5JUU;[69]). This disordering occurs in proximity to the sarcin-ricin loop (SRL), a crucial ribosomal site for triggering GTP hydrolysis. Post-hydrolysis, domains III and V shift relative to Switch 2 in domain I, with domain III rotating ∼5 degrees and domains IV and V rotating together by ∼9 degrees. This movement repositions the diphthamide-containing tip of domain IV by ∼6 Å toward the top of helix 44 in the small ribosomal subunit (2P8X; [38], 5JUU; [69]).

In our structures, eEF2 exhibits hallmark features of the post-GTP hydrolysis state: The Switch I loop is partially disordered; although the backbone could be modeled in both classes, the His108 side chain is only faintly visible in the IRES-containing eEF2 complex and better visible in the canonical eEF2 complex. Switch II is completely unresolvable in both, which is consistent with a GDP-bound or nucleotide-release state. Additionally, domain II is rotated relative to the GTP-bound conformation, and the diphthamide-containing loop of domain IV is retracted by approximately 6 angstroms (Figure 3F). While no nucleotide density is observed in the IRES-bound eEF2 class, the canonical eEF2 class contains interpretable density for the nucleoside and two phosphates. We attempted to model both GTP and GDP into the density, and found that GDP provides a better fit (Figure S8A). These features clearly distinguish our structures from previously published GTP-bound eEF2 conformations such as 6GZ5 [40], where both Switch loops were resolved. Together, the data support the assignment of our eEF2 conformations to the post-GTP hydrolysis state.

Domain IV of eEF2 contains three loops – loop 1, loop 2, and loop 3, all critical for its function. Consistent with the results of Flis et al. (2018) [40] and Djumagulov et al. (2021) [42], our analysis shows that the diphthamide residue of domain IV stacks against the ribose of A36 in peptidyl-tRNA (Figure S8B). Loop 2 interacts with helix 32, anchoring domain IV near the A site, while loop 3 engages with 18S rRNA h44, likely to stabilize the reading frame (Figure S8B). Additional stabilizing interactions, previously underexplored, occur between eEF2 domain IV and 28S rRNA of the 60S subunit, involving water- or metal-ion-mediated contacts: Trp 685 with C1’ of C3761 (7.22 Å), Ile 722 with C1’ of A3760 (5.58 Å), and Gly 719 with C1’ of A3760 (5.1 Å) (Figure S8C, S8D).

Overall, the comparison of these two structural classes (one with IRES bound 80S in complex with eEF2 and the other is canonical mRNA containing eEF2) highlights a clear transition from an IRES-dependent initiation complex to a canonical elongation complex (Figure S9), with eEF2 playing a central role in coordinating tRNA translocation and stabilizing the ribosomal complex. While the first class still retains a partially visible IRES, the second class represents a fully canonical elongation state with only the ORF mRNA present along with hybrid tRNAs bound to the 40S subunit. Both classes feature a slightly rotated ribosome.

### The structure of an eEF1A-bound post-proofreading complex

One of the classes from our elongation setup, which includes eEF1H, eEF2 and tRNA^Ala^, shows a clear map coverage for eEF1A, A*/T-Ala-tRNA^Ala^, P-tRNA^Ala^, E-tRNA^Ala^, and the translocated mRNA encoding Ala-Thr-His-Stop (Figure 4A – 4C). Determining the exact codon positions is complicated by the blurred density of mRNA due to residual heterogeneity within the class. Despite these difficulties, the assured absence of any tRNAs other than tRNA^Ala^ allowed us to place the Ala codon (GCU) in the P site and model the mRNA from there as part of a singly-translocated complex. In this complex, the A site contains a mismatched codon-anticodon pair, resulting in a disordered anticodon stem but a well-resolved rest of the tRNA, indicative of an Ala-tRNA^Ala^ in the process of being rejected from the A site (Figure 4A – 4D). Furthermore, the structure of eEF1A shows significant differences when compared to published structures, with an RMSD of 1.28 Å compared to the structure of ribosome-bound eEF1A reported by Shao et al. (PDB ID 5LZS [47]). Our eEF1A is shifted approximately 12 Å away from the A site (Figure 4D, right panel, S10A). Comparison of our A*/T-tRNA with the A/T-tRNA in 5LZS [47] revealed a very similar conformation, except for the highly flexible residues 31–39, which could not be modeled (Figure 4D, left panel). The acceptor arm of our A/T-tRNA positions G1 near eEF1A residues Y254 and R322 (Figure S10C), while A76 nestles into a hydrophobic pocket formed by I259, G258, G257, I256, and Y254 (Figure S10D), resulting in tight binding of the tRNA to eEF1A.

**Figure 4.**
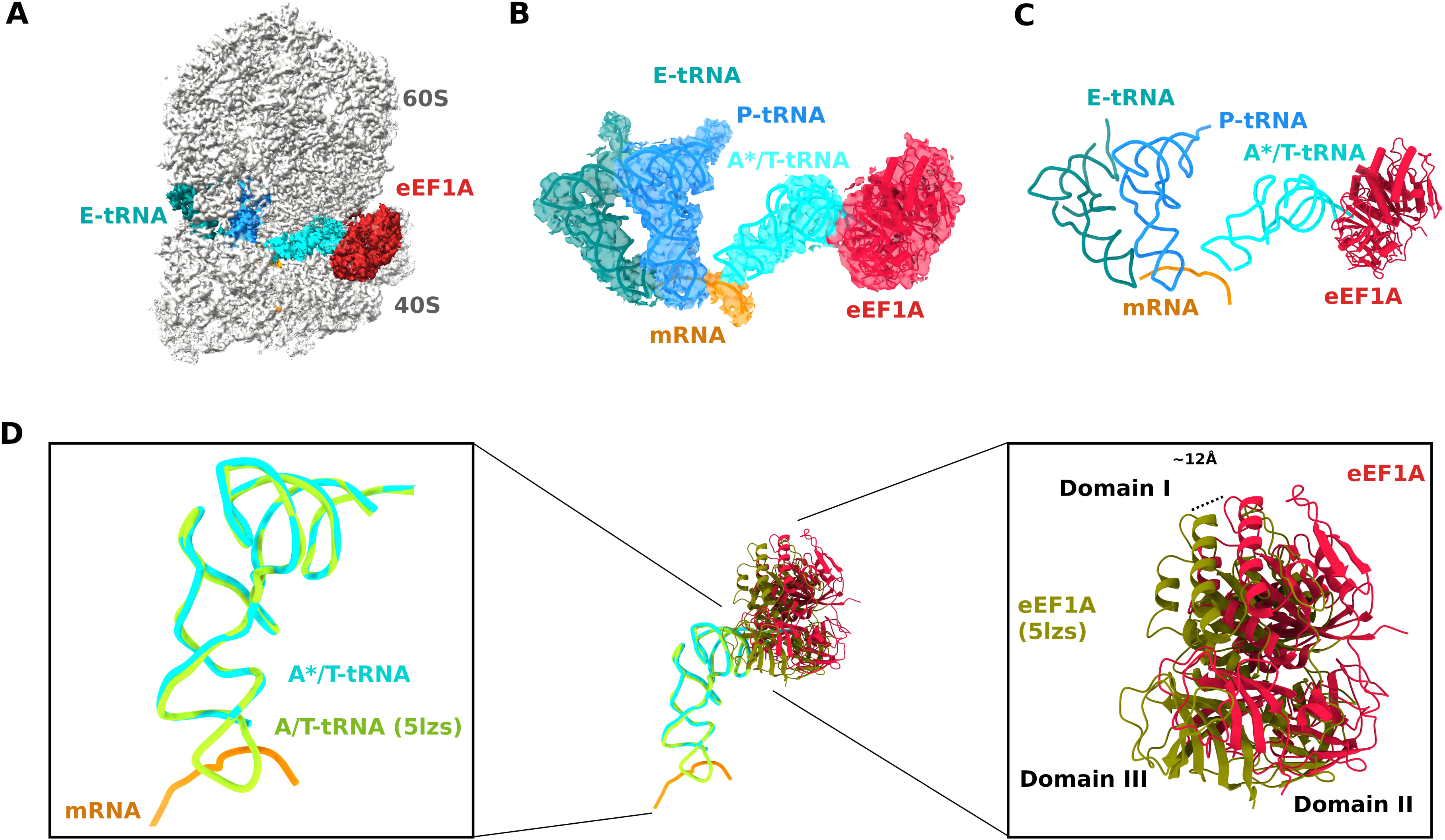
eEF1A-Mediated tRNA Delivery and Proofreading. (A) Cryo-EM map of a canonical A*/T complex, featuring an E-site tRNA (teal), a P-site tRNA (dodger blue), and an A*/T-tRNA (cyan) bound to eEF1A (crimson). This structure captures a key intermediate state during the proofreading process, where a mismatched tRNA is being rejected from the A site. (B) and (C) Focused views of the complex in (A), showing the cryo-EM map (B) and the atomic model (C). These panels highlight the positioning of eEF1A and the A*/T-tRNA, demonstrating the unique proofreading conformation where the eEF1A is shifted along its axis away from the A site. (D) A direct structural comparison of our proofreading complex with the Didemnin B-trapped, cognate complex (PDB ID 5LZS). The right panel highlights the significant ∼12 Å shift of eEF1A away from the 28S rRNA compared to the 5LZS model, reflecting the conformational changes associated with the rejection of a mismatched tRNA.

It is important to point out that the 5LZS complex [47] was trapped with Didemnin B and a fully paired codon-anticodon pair, whereas our structure was captured in the presence of GTP and an active elongation complex with a mis-paired codon-anticodon pair. Compared with the 5LZS model [47], which features an arrested eEF1A·A/T-tRNA complex, eIF1A in our structure has shifted away from the 28S residue 4605 by ∼ 12Å, accompanied by a slight rotation of the entire domain away from the A*/T tRNA. Additionally, we find the anticodon loop and adjacent parts of the tRNA to be distorted, with the A44 residue displaced by ∼12Å compared to the conformation seen in cognate base pairing [47] (Figure 4D). In our structure, the A*/T-tRNA interacts with uL11 (C57 with Thr 25), and eS30 (backbone interactions between residues 25-26 and amino acids 76-80) which stabilize its position in the A site (Figure S11A, S11B). Concurrently, eEF1A is tightly bound to the A*/T-tRNA and engages in multiple direct interactions between its domain III ɑ-helices and the tRNA backbone (Figure S10C, S10D). Additionally, eEF1A interacts with 28S rRNA through domain I and 18S rRNA through domain II, contributing to the stabilization of the mismatched TC in the A site. Furthermore, we observed a direct interaction of eEF1A domain II with uS12 Arg142, potentially aiding in positioning the eEF1A.tRNA complex into the A-site cleft (Figure S10B). The P- and E-site tRNAs are in canonical conformations with the ribosome in an almost non-rotated position, with a 1.4-degree rotation of the 40S body.

## DISCUSSION

Initiation of translation mediated by the NediV IRES involves its direct binding to the P site of the 80S ribosome, and this initiation process is therefore even simpler than on type 6a and type 6b IRESs, which involves binding of the IRES to the A site followed by pseudo-translocation to the P site. Here, we have determined the structures of ribosomal complexes at key stages during initiation and early elongation on this IRES.

### Initiation on the NediV IRES

The multiple stabilizing interactions of the IRES with the ribosome in binary complexes (Figure 1A, S2) likely account for its ability to bind directly to the P site, bypassing the pseudo-translocation step, and for the strong preference of this class of IRES for binding to 80S ribosomes rather than to 40S subunits. The ability of the IRES to maintain a stable interaction with the ribosomal P site in the 80S·NediV IRES·aa-tRNA complex, even across varying rotational states (Figure 2A), is instrumental in facilitating initiation. This complex stabilizes the bound IRES and its interactions with the ribosome, places the IRES-linked ORF into the mRNA-binding channel and positions the tRNA into the A site for subsequent elongation steps. It therefore represents a crucial intermediate in this translation mechanism.

Destabilization of PKI may begin during or immediately after the initial translocation step (Figure 2H, S9), similar to the disassembly of the analogous domain in the CrPV IRES upon its transition from P to E sites [29]. The presence of the destabilized PKI in the E site of the 80S·IRES^E^·aa-tRNA^P^·eEF2 complex and the absence of E-site tRNA (Figure 2C, 2F) might account for two characteristics of IGR IRES-mediated translation. Failure to establish an interaction between the 3′-terminal adenosine (A76) of E-site tRNA and 23S rRNA nucleotide C2394 in bacterial ribosomes decreases the rate of translocation [78,79]. A similar phenomenon could account for the slow rate of early cycles of translocation observed after initiation on IGR IRESs [19,80], when the E site is either vacant or contains the destabilized PKI, and has not yet been replaced by deacylated tRNA. Premature release of E-site tRNA from bacterial ribosomes is associated with high-level frameshifting [81] and analogously, the absence of base-pairing with deacylated tRNA in the E site at this early stage might contribute to the frameshifting observed during initiation on NediV and other IGR IRESs [34–36,82].

The movement of the A-site tRNA to the P site during the first translocation cycle (Figure 4B, 2C, S9) has the potential to introduce steric clashes with IRES PKI, initiating its melting, and thereby facilitating its clearance from the E site, triggering a cascade that ultimately leads to dissociation of IRES domains 2 and 3 from the ribosome. Destabilization of PKI may thus facilitate the clearance of the E site by the IRES and may also aid in threading the viral ORF2 mRNA into the mRNA channel, ensuring a smooth transition to elongation. Given the observed rigidity of IRES interactions with the ribosome before translocation, this transition may represent a tightly regulated step that balances IRES-mediated control with the canonical elongation mechanism. The presence of eEF2 in a post-GTP hydrolysis state (Figure 3F) further suggests that this process is coordinated with the conformational dynamics of elongation factors, reinforcing the idea that once translocation occurs, the IRES relinquishes its structured hold on the ribosome, enabling productive translation of the viral message.

### Early steps in elongation

The capture of the singly translocated 80S·IRES^E^·aa-tRNA^P^·eEF2 complex, featuring eEF2 bound to the ribosome with tRNA occupying the P site, offers a crucial snapshot of this transition from IRES-mediated initiation to canonical elongation. The observed near-zero-degree rotation of the ribosome, coupled with the precise positioning of eEF2, suggests a highly orchestrated handoff. This state likely represents a late-stage translocation intermediate, poised for eEF2 release in which the ribosome is primed for the commencement of canonical elongation. It is plausible that eEF2 stabilizes this specific conformation, ensuring an efficient and seamless transition. A key structural feature of this complex is the ∼6°rotation in IRES domains 2 and 3 towards the 40S head (Figure 2G – 2I), accompanied by the destabilization of the PKI domain in the E site.

The close proximity of the P-site and E-site tRNAs within our canonical 80S·tRNA^E^·aa-tRNA^P^·eEF2 complex (Figure S7) raises intriguing questions about the dynamics of tRNA release. The electrostatic repulsion between the closely packed tRNAs in P and E sites may be utilized for promoting E-site tRNA release. It could serve to ensure efficient ribosome clearance before the next elongation cycle, preventing potential clashes and maintaining translational fidelity. The position of the P gate observed, which prevents full tRNA accommodation, might represent a paused state, allowing quality control or coordination with other cellular processes.

While the precise chemical state of the nucleotide on eEF2 remains unresolved due to resolution limitations, the observed density for eEF2 aligns with expected post-GTP hydrolysis conformations. This highlights the conserved role of the Switch 1 loop in transmitting conformational signals within eEF2 and coordinating ribosome-elongation factor interactions [38,41]. The presence of eEF2 in this state further implies that these interactions stabilize a translocation-ready conformation, setting the stage for elongation factor release and continued mRNA-tRNA movement. Finally, the critical role of the diphthamide modification in eEF2 in maintaining reading-frame integrity, as suggested by previous studies [40–42], is supported by our structural data. The observed interactions (Figure S8B) suggest that in this complex, diphthamide may play a crucial role in preventing mRNA backsliding during P-gate disruption as tRNAs and mRNA transition. Collectively, these findings underscore the central role of eEF2 in ribosomal translocation, by pointing to the existence of a highly conserved mechanism across IRES-mediated and canonical elongation pathways.

### A post-proofreading state of the eEF1A-bound ribosome after GTP hydrolysis

The error rate of translation in eukaryotes is ∼10^-3^ to 10^-5^ per codon [83,84], a level of accuracy that cannot be accounted for solely by the difference in binding energy of cognate and near-cognate tRNAs to the A site codon [85,86]. Instead, accuracy results from two-step kinetic proof-reading [87]. During initial selection, EF-Tu/eEF1A·GTP·aa-tRNA samples the A site codon of mRNA, and establishment of codon-anticodon base-pairing induces conformational changes in the ribosome that trigger GTP hydrolysis [88]. The aa-tRNA is released from eEF1A·GDP and accommodated in the PTC, followed by peptide bond formation. Fidelity results from rejection of incorrect aa-tRNAs at the initial selection stage and during subsequent proofreading.

The presence of only one type of tRNA (tRNA^Ala^) in the reaction mixture and a mismatched codon at the A site have enabled us to capture a complex representing a key step of the tRNA rejection process. The persistence of this complex is consistent with changes in the ribosome and ternary complex, which makes proofreading a rate-limiting step in elongation [72], and because rejection can occur multiple times with the same mismatched tRNA, a significant number of ribosomes are trapped in this state. The structure of the eEF1A-bound complex presents a compelling snapshot of a post-proofreading state (Figure 4), deviating significantly from previously captured complexes with cognate tRNA [47,72,73]. Notably, the eEF1A-containing structure described here is the first example of this complex visualized in the presence of GTP without any trapping agents, as well as with a single species of mismatched tRNA. This result highlights the advantage of using IGR IRESs for the structural characterization of specific steps in eukaryotic translation in defined conditions. The substantial shift of eEF1A and the A/T-tRNA away from the A site, coupled with the distorted anticodon loop, strongly suggests an active rejection mechanism. The observed 1.4-degree rotation of the 40S body could also play a role in this process, potentially contributing to the destabilization of the mismatched codon-anticodon interaction. The extensive interactions between A*/T-tRNA and the ribosome, including uL11, rRNA, and eS30 (Figure S6A, S6B), might serve to maintain the rejected tRNA in a holding pattern, prior to its eventual release. It is possible that the interactions with uS12 Arg142 (Figure S10B) is a key component to the proofreading mechanism, as the same interaction is not seen in previous structures. Future studies employing time-resolved cryo-EM or biochemical assays could further elucidate the precise sequence of events leading to tRNA rejection and provide a more complete understanding of eEF1A’s role in maintaining translational fidelity.

This study provides key structural insights into the mechanism of translation initiation and early elongation mediated by the NediV IRES. We have shown how the NediV IRES directly binds to the ribosomal P site and how its stable interactions facilitate the positioning of the viral ORF for translation. Furthermore, our findings illuminate the dynamic transitions during early elongation, particularly the role of eEF2 and eEF1A in coordinating the handoff from IRES-mediated initiation to canonical elongation and the process of tRNA rejection during proofreading. These structures, particularly the unprecedented post-proofreading state captured without trapping agents, offer a deeper understanding of fundamental eukaryotic translation mechanisms and highlight the utility of IGR IRESs in dissecting these complex processes.

### Depositions

All cryo-EM maps and corresponding atomic models from this study have been deposited in the EMDB and PDB. The P-site complex is available under accession numbers PDB ID 9Q1S and EMDB 72137, and the A-site complex under PDB ID 9Q1Q and EMDB 72136. The A-tRNA complex is deposited as PDB ID 9Q2O and EMDB 72170, the canonical eEF1A complex as PDB ID 9Q2P and EMDB 72171, the canonical eEF2 complex as PDB ID 9Q2M and EMDB 72168, and the IRES–eEF2 complex as PDB ID 9Q2T and EMDB 72175.

## Acknowledgments

This work was supported by grants from the National Institutes of Health: R01 GM29169 and R35 GM139453 to J.F., R35 GM122602 to T.V.P., and R01 GM097014 and R21 AI188505 to C.H. We want to thank Thea Hu (University of Bristol) for her help in PDB curation, proofreading, and preparing the graphical abstract.

## Authors contributions

SD, CA, PD, FAR, PG, and ZB analyzed the data. IA and CA were responsible for sample preparation and data collection. The manuscript was written by SD, TP, CH, and JF. TVP, CH, and JF supervised the project.

## Supplementary Figure Legends

**Figure S1. Close-up Views of Key Structural Motifs within the NediV IRES**

Detailed views of the NediV IRES, highlighting the conserved pseudoknot domains (PKI, PKII, PKIII), the variable region loop (VRL), and the helices (P3.1 and P3.2) that define its characteristic architecture.

**Figure S2. Ribosomal Contacts of the NediV IRES (P-site)**

(A) A global view of the P-site-bound IRES and its extensive network of interactions with the 80S ribosome, showcasing the points of contact between the viral RNA and both ribosomal subunits. (B) and (C) Focused views of the IRES-ribosome contacts, showing the cryo-EM maps

(B) and atomic models (C) to illustrate specific stabilizing interactions. (D) A detailed view of the interaction between IRES Domain 1 and the ribosomal L1 stalk, which acts as a key anchor point for the IRES. (E) Close-up view of the contact between the highly conserved L2.2 loop of the IRES and ribosomal protein eS25 at the head of the 40S subunit. (F) Additional stabilizing contacts between the IRES and ribosomal proteins uS4, uS5, and eS30.

**Figure S3.** (A) Superposition of the rotated (blue) and non-rotated (purple) 60S subunits within the 80S ribosomes bound to the NediV IRES, illustrating the subtle rotational dynamics observed in the complexes. (B) A top-down view of the 40S and 60S subunits. (C) A focused view of the arrangement of expansion segment ES7L in the rotated complex. (D) The full ribo-nucleotide sequence of the NediV IRES and the viral open reading frame (ORF) annotated in green.

**Figure S4. Detailed IRES-Ribosome Interactions**

(A) A close-up view of the interactions between the IRES and the A-gate residues (nt. 1701 and nt. 1248) of the 18S rRNA, which are critical for stabilizing the IRES in the P site. (B) Detailed view of the IRES’s interaction with ribosomal protein uS11. (C) Detailed view of the IRES’s interaction with ribosomal protein uS5.

**Figure S5.** A comparison of the orientation of G626 in our A-tRNA-bound IRES complex relative to the published structure (PDB ID 5LZS).

**Figure S6.** (A) A global view showing the path of the viral open reading frame (ORF) mRNA (dark orange) through the mRNA cleft, following translocation of the IRES. (B) A local view showing the detailed interactions of the ORF mRNA with the 18S rRNA (residues 1825 and 1701).

**Figure S7.** A close-up view highlighting the unusually short distance between the backbones of the P-site and E-site tRNAs in the canonical elongation complex, suggesting a unique structural arrangement.

**Figure S8. eEF2 Conformational Dynamics**

(A) A comparison of the Switch I and Switch II loops in eEF2 from our canonical complex with the GTP-bound structure (PDB ID 6GZ5). This figure highlights the disorder in our structure, consistent with a post-GTP hydrolysis state. (B) Position of the diphthamide residue and the critical loops 1, 2, and 3 of eEF2 Domain IV relative to the tRNA, mRNA, and 18S rRNA. (C) A schematic of the major contact sites between eEF2 and the 28S rRNA of the 60S subunit. (D) A detailed view of the contacts between eEF2 residues (Trp 685, Ile 722, Gly 719) and the 28S rRNA (C3760 and C3761).

**Figure S9. Schematic of NediV IRES-mediated translation initiation process**

A schematic illustration summarizing the NediV IRES-mediated translation initiation process, the transition to the first elongation step, and the key conformational changes that occur.The question mark next to the arrow from the A-site loaded state to the P-site loaded state indicates that this transition has not been experimentally established.

**Figure S10. eEF1A Domain Structure and Interactions**

(A) An overview of the eEF1A domain structure with the cryo-EM density map, showing the three distinct domains. (B) A detailed view of the interactions between eEF1A and ribosomal protein uS12 (Arg 142) and the A*/T tRNA, which helps position the complex in the A-site cleft. A close-up of the interaction between eEF1A residues R322 and Y254 and the G1 residue of the A*/T tRNA. (D) A focused view of the hydrophobic pocket formed by eEF1A residues that tightly binds the A77 residue of the A*/T tRNA.

**Figure S11. A/T tRNA Stabilization**

(A) A detailed view of the stabilizing interactions between the A*/T tRNA and ribosomal protein eS30. (B) Detailed view of the interactions between the A*/T tRNA and ribosomal protein uL11.

**Table 1: Cryo-EM and Refinement Statistics**

**Table 2: Cryo-EM workflow**

