## Supplementary material for "Structural Studies of Nedicistrovirus IRES-Driven, Initiation Factor-independent Translation Shed Light on Key Steps of Eukaryotic Translation Elongation": Table 2

### Table2

Images

1.Motion Correction
2. CTF estimation
3. Gautomatch

1. Particle Extraction
2. 2D Classification
3. Select all 80S particles

3D Refinement

CryoDRGN
heterogeneous reconstruction

Selected
Classes

NediV-80S (P-site)
PDB: 9Q1S
EMDB: 72137

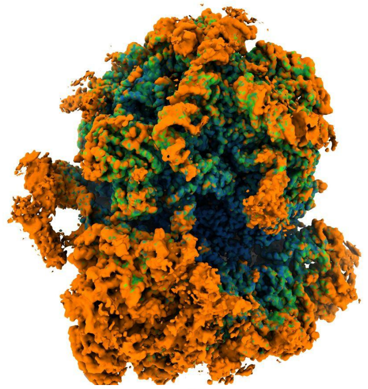

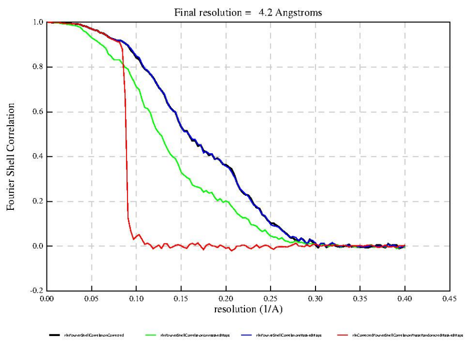


NediV-80S (A-site)
PDB: 9Q1Q
EMDB: 72136

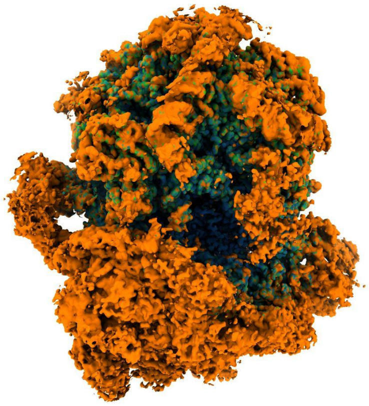

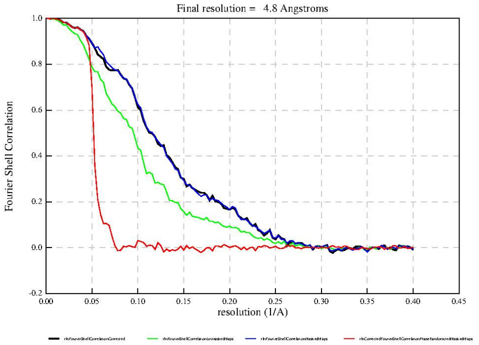


NediV-80S-A-site tRNA
PDB: 9Q2O
EMDB: 72170

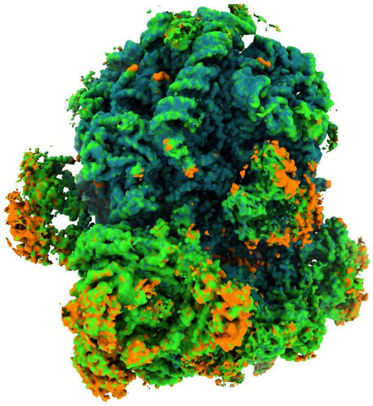

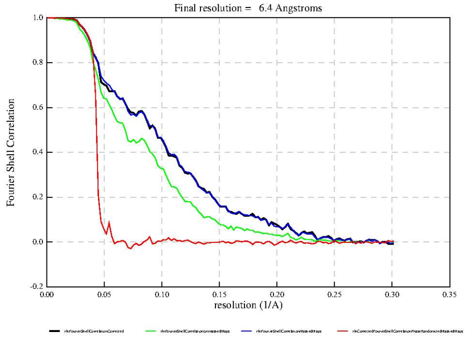


NediV-80S-P-site tRNA-eEF2
PDB: 9Q2T

EMDB: 72175

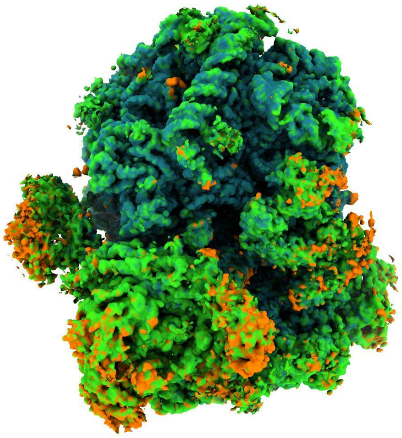

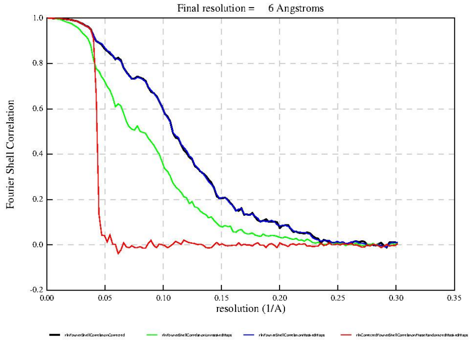


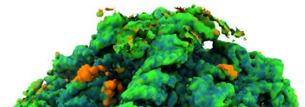

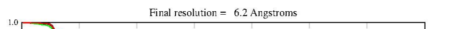
