## Supplementary material for "Structural Studies of Nedicistrovirus IRES-Driven, Initiation Factor-independent Translation Shed Light on Key Steps of Eukaryotic Translation Elongation": Table 1

**Cryo-EM data collection, refinement and validation statistics**

| Ribosomal complex | P-site IRES (PDB 9q1s EMDB 7137) | A-site IRES (PDB 9q1q EMDB 72136) | P-site IRES A-site tRNA (PDB 9q2o EMDB 72170) | eEF2 IRES (PDB 9q2t EMDB 72175) | canonical eEF1A (PDB 9q2p EMDB 72171) | canonical eEF2 (PDB 9q2m EMDB 72168) |
|---|---|---|---|---|---|---|
| **Data collection and processing** | | | | | | |
| Electron Microscope | Polara F30 | Polara F30 | Polara F30 | Polara F30 | Polara F30 | Polara F30 |
| Detector | K2 Summit | K2 Summit | K2 Summit | K2 Summit | K2 Summit | K3 Summit |
| Magnification | 31,000 | 31,000 | 31,000 | 23,000 | 23,000 | 39,000 |
| Voltage (kV) | 300 | 300 | 300 | 300 | 300 | |
| Electron exposure (e–/Å$^2$) | 42 | 42 | 23 | 23 | 23 | 71 |
| Defocus range (μm) | -3.5 to -1.5 | -3.5 to -1.5 | -3.5 to -1.5 | -3.5 to -1.5 | -3.5 to -1.5 | -3.5 to -1.5 |
| Pixel size (Å) | 1.24 | 1.24 | 1.66 | 1.66 | 1.66 | 0.95 |
| Symmetry imposed | | | | | | |
| Initial particle | C1 | C1 | C1 | C1 | C1 | C1 |
| images (no.) | 459177 | 459177 | 546318 | 546318 | 546318 | 220763 |
| Final particle images (no.) | 48932 | 17777 | 38447 | 14021 | 38447 | 20663 |
| Map resolution (Å) FSC threshold | 0.143 | 0.143 | 0.143 | 0.143 | 0.143 | 0.143 |
| Map resolution range (Å) | 4.2 | 4.8 | 6.4 | 5.9 | 6.1 | 5.1 |
| **Refinement** | | | | | | |
| Initial model used (PDB code) | 5LZS | 5LZS | 5LZS | 6GZ5 | 5LZS | 6GZ5 |
| Model composition | | | | | | |
| Atoms | 379106 | 378361 | 381691 | 393781 | 385964 | 393441 |
| Protein residues | 11855 | 11862 | 11877 | 12482 | 12153 | 12699 |
| Nucleotide residues | 5681 | 5652 | 5749 | 5830 | 5748 | 5706 |
| Ligands | Zn:8 Mg:275 | Zn:8 Mg:275 | Zn:8 Mg:275 | Zn:5 Mg:326 | Zn:8 Mg:276 | Zn:5 Mg:326 GDP:1 |
| R.m.s. deviations | | | | | | |
| Bond lengths (Å) | 0.004 | 0.004 | 0.003 | 0.005 | 0.003 | 0.003 |
| Bond angles (°) | 0.829 | 0.793 | 0.627 | 0.689 | 0.640 | 0.675 |
| Validation | | | | | | |
| MolProbity score | 1.77 | 1.74 | 1.87 | 1.68 | 1.84 | 1.69 |
| Clashscore | 4.63 | 4.30 | 4.89 | 1.46 | 3.15 | 1.46 |
| Poor rotamers (%) | 1.94 | 2.03 | 3.03 | 2.50 | 5.04 | 2.59 |
| Ramachandran plot | | | | | | |
| Favored (%) | 95.43 | 95.77 | 96.25 | 91.62 | 96.81 | 91.80 |
| Allowed (%) | 4.48 | 4.20 | 3.74 | 8.30 | 3.15 | 8.14 |
| Disallowed (%) | 0.09 | 0.03 | 0.01 | 0.07 | 0.04 | 0.06 |
