## Supplemental Figures for "Structural Studies of Nedicistrovirus IRES-Driven, Initiation Factor-independent Translation Shed Light on Key Steps of Eukaryotic Translation Elongation"

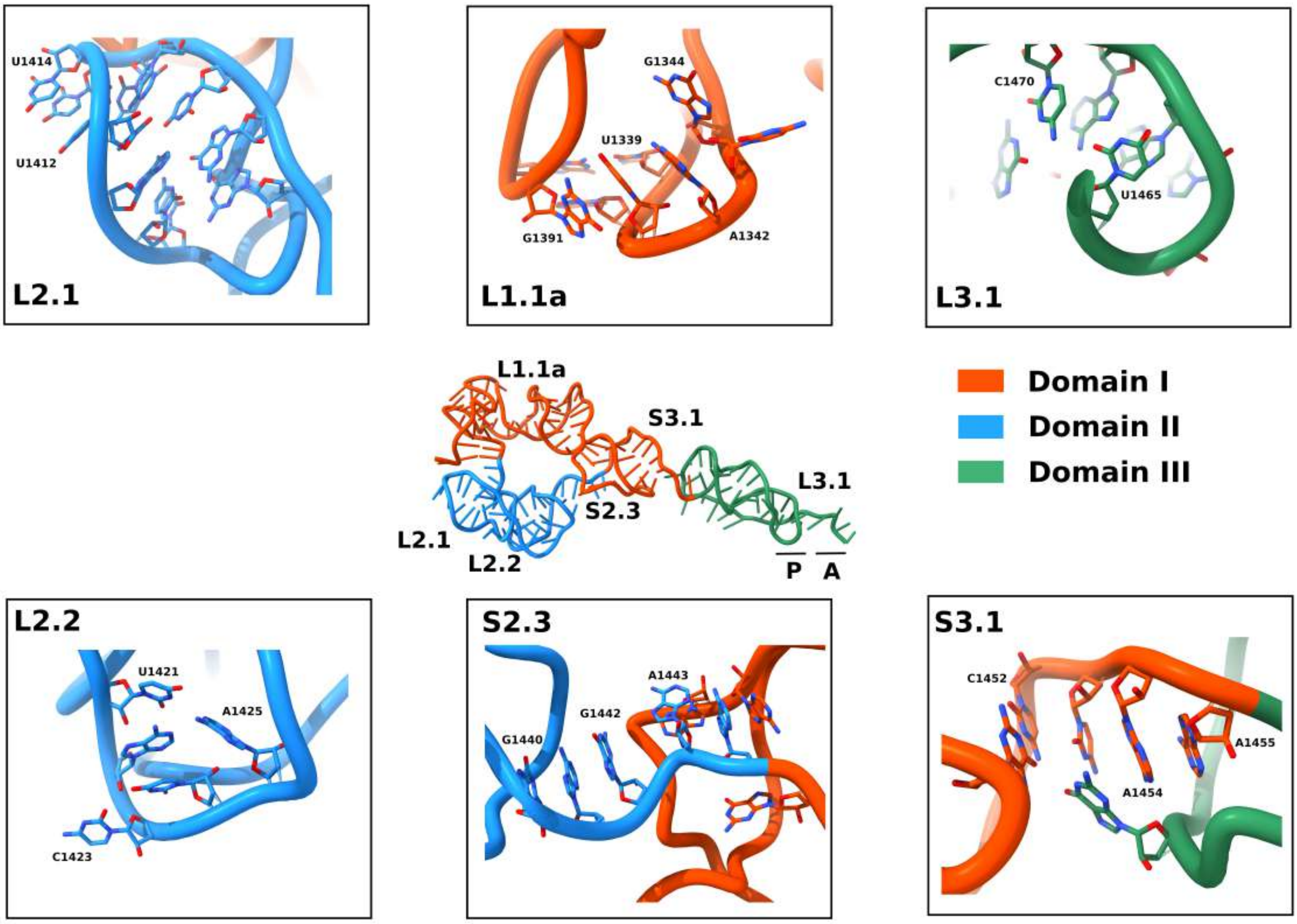
U1414
U1412
L2.1
G1344
U1339
G1391
A1342
L1.1a
C1470
U1465
L3.1
L1.1a
S3.1
L3.1
S2.3
L2.1
L2.2
P
A
Domain I
Domain II
Domain III
L2.2
U1421
A1425
C1423
S2.3
A1443
G1442
G1440
S3.1
C1452
A1455
A1454

**Figure S1**

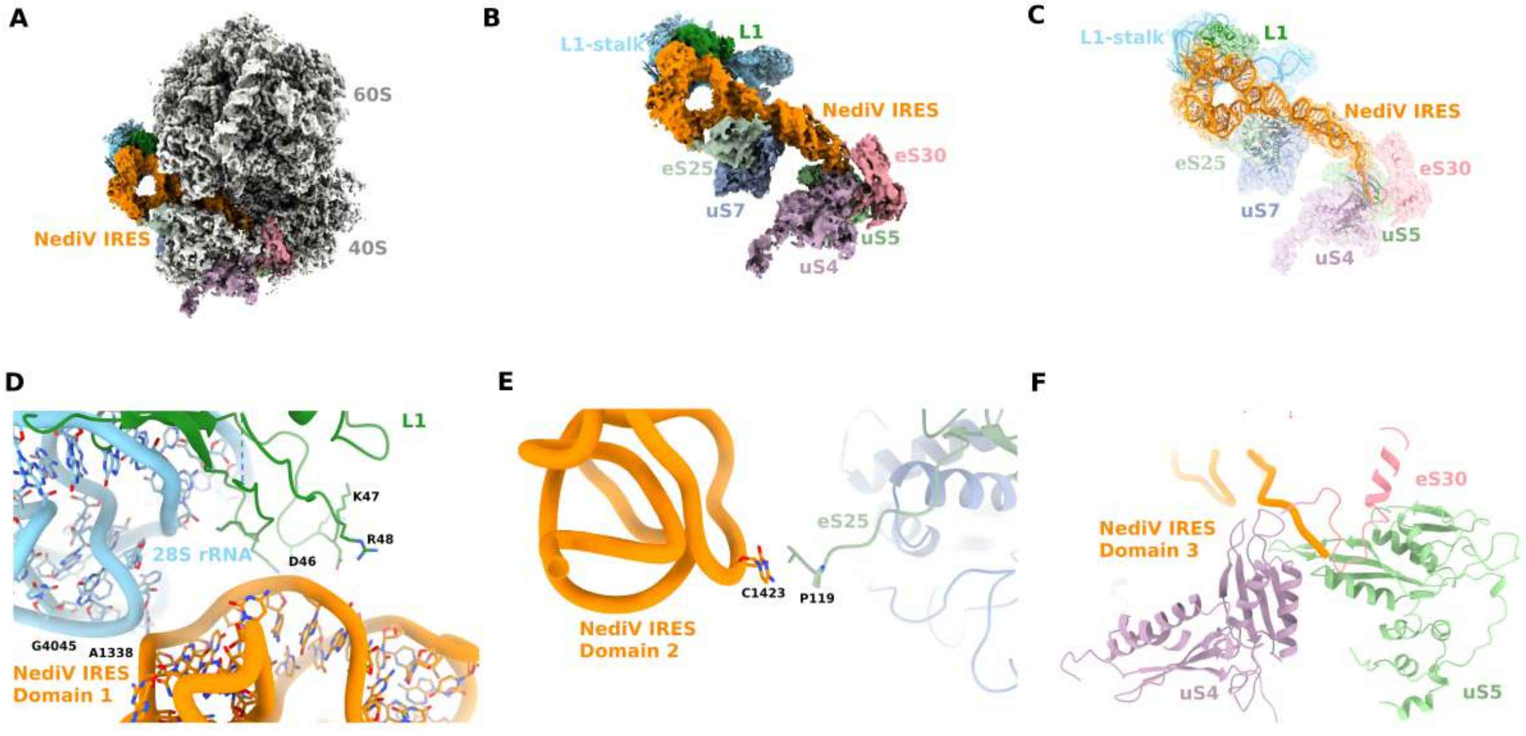
A
60S
NediV IRES
40S
B
L1-stalk
L1
NediV IRES
eS25
eS30
uS7
uS5
uS4
C
L1-stalk
L1
NediV IRES
eS25
eS30
uS7
uS5
uS4
D
L1
K47
R48
28S rRNA
D46
G4045
A1338
NediV IRES
Domain 1
E
eS25
C1423
P119
NediV IRES
Domain 2
F
eS30
NediV IRES
Domain 3
uS4
uS5

### Figure S2

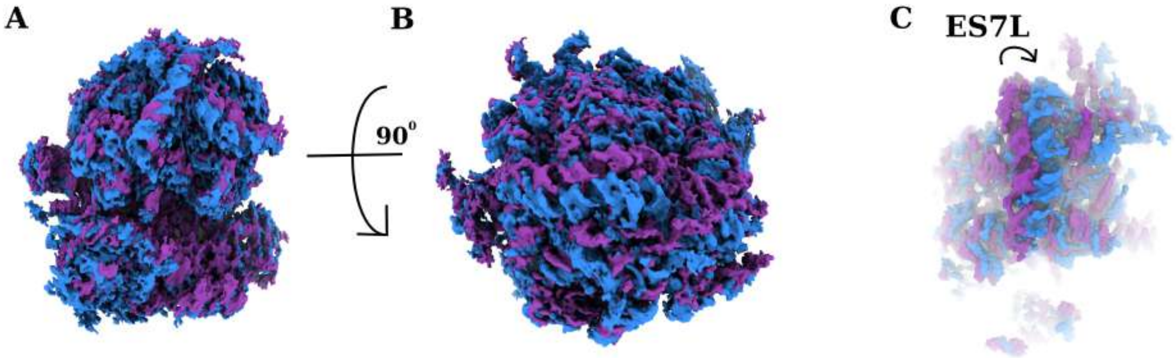


D

GGAUCCUAAUACGACUCACUAUAGGCCGACCCGGUGACGGGUCGGCCUAAGUU
GGAUGGCACACUUGGUGAGAUGUGCCUCCAUCCCCAGGAGUUUUGGGACGAGC
AUGCACCUAAAGUUAUGGCCGUUAUGCGUGAGCAUUUGGGUGUUGAACCCAUG
UUUACCUCACGCGAGGGCUAUCUUGAGGAGACAGCUGCCCGCACCGACUUCUG
GUUUUAGGUUGUAUAUACGGUGCUCUUUUAGUAAAUAAUUGGGAUUCCCCCUAU
GGAGCAUGACAGGUAAGCCAUUUCACCCCCCAUUUUCAUGGUACUACUCAGGUU
GGACCAGAGAAGGAUUCCCCAAGGCUGCCUUUGGGACAAAGCAGCUUGUAUAUA
GUCCCGCUACACAUAAUAAUGAUAGUAUAGAGAGAGAUACGAUCUCUGAGGAAG
UCAAUCCUGGAUUGUCAGUUCAAGGCAAUCACACCAUUGGUUUAACCACUUUUC
CUGUUGAGUCCUCUAGA

**Figure S3**

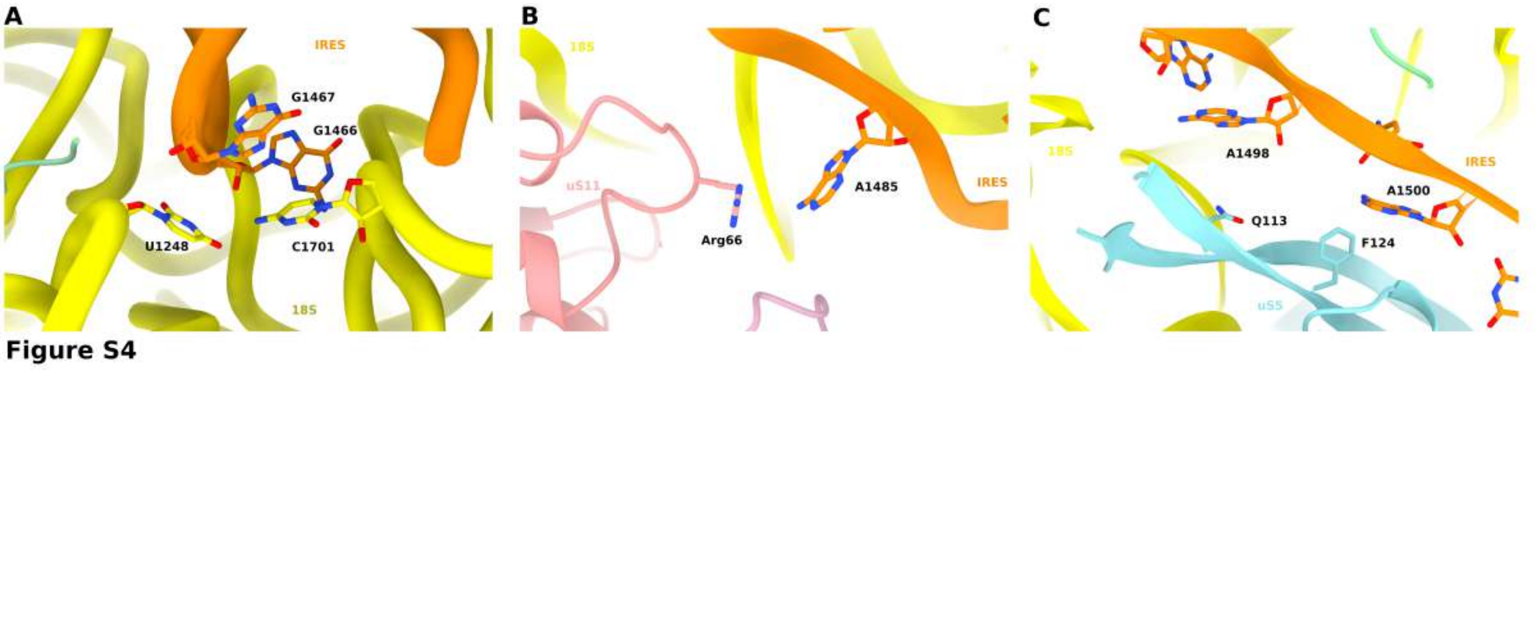
A
IRES
G1467
G1466
U1248
C1701
18S
B
18S
uS11
Arg66
A1485
IRES
C
18S
A1498
IRES
A1500
Q113
F124
uS5
Figure S4

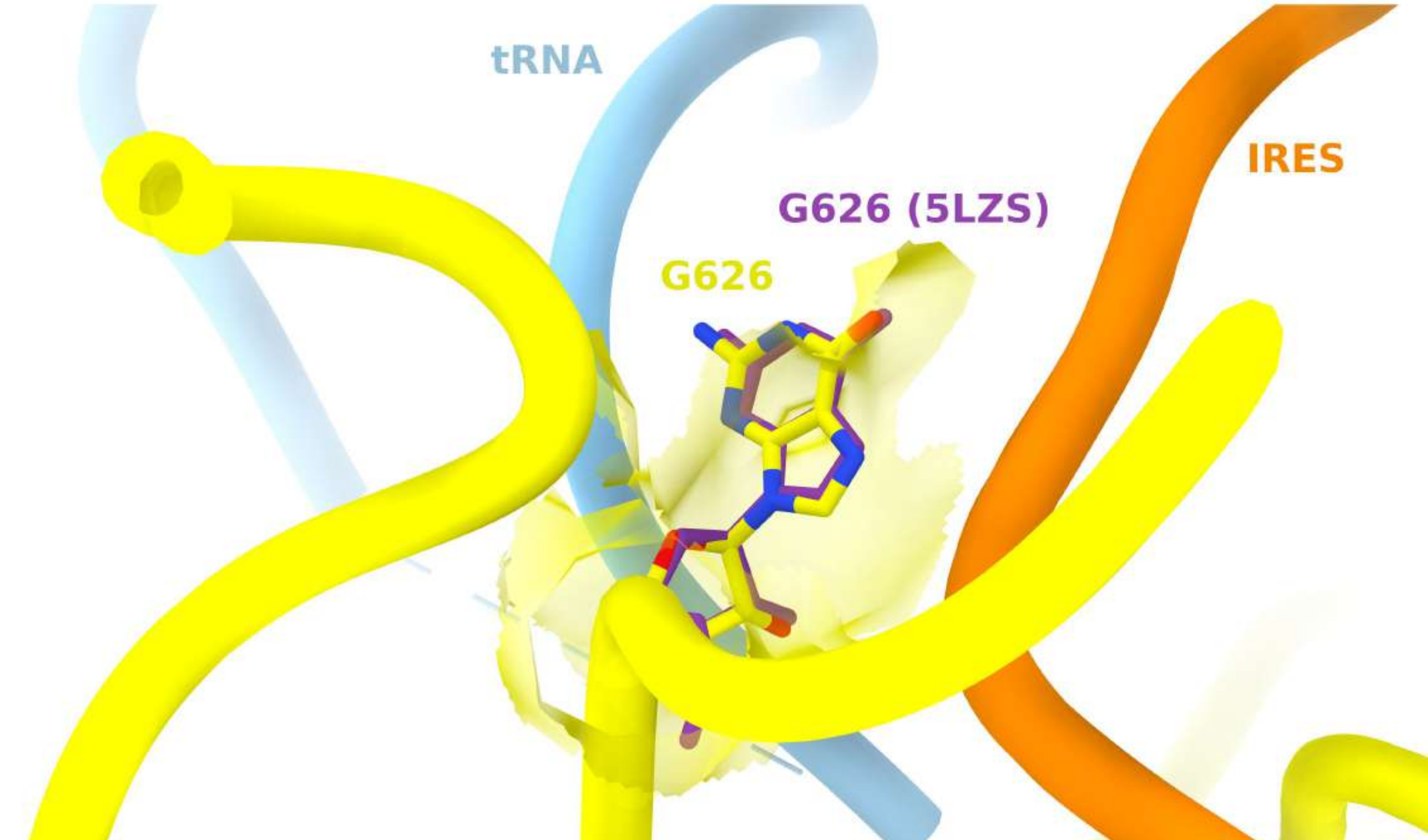


### Figure S5

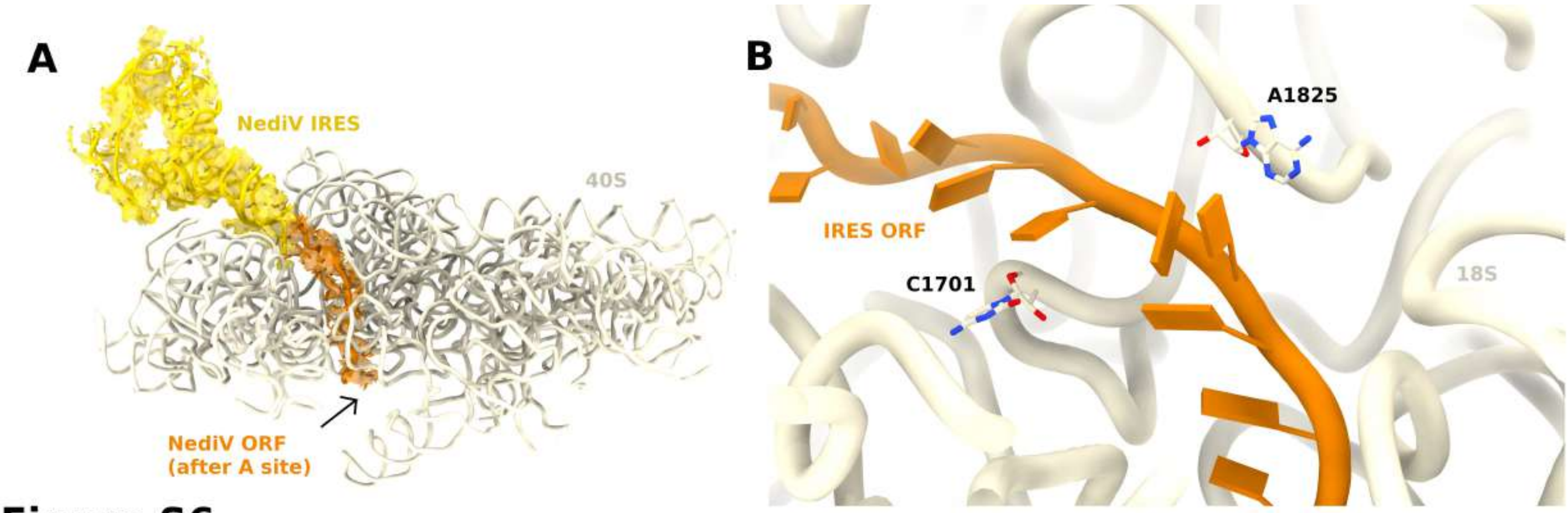
A
NediV IRES
40S
NediV ORF
(after A site)
B
A1825
IRES ORF
C1701
18S

**Figure S6**

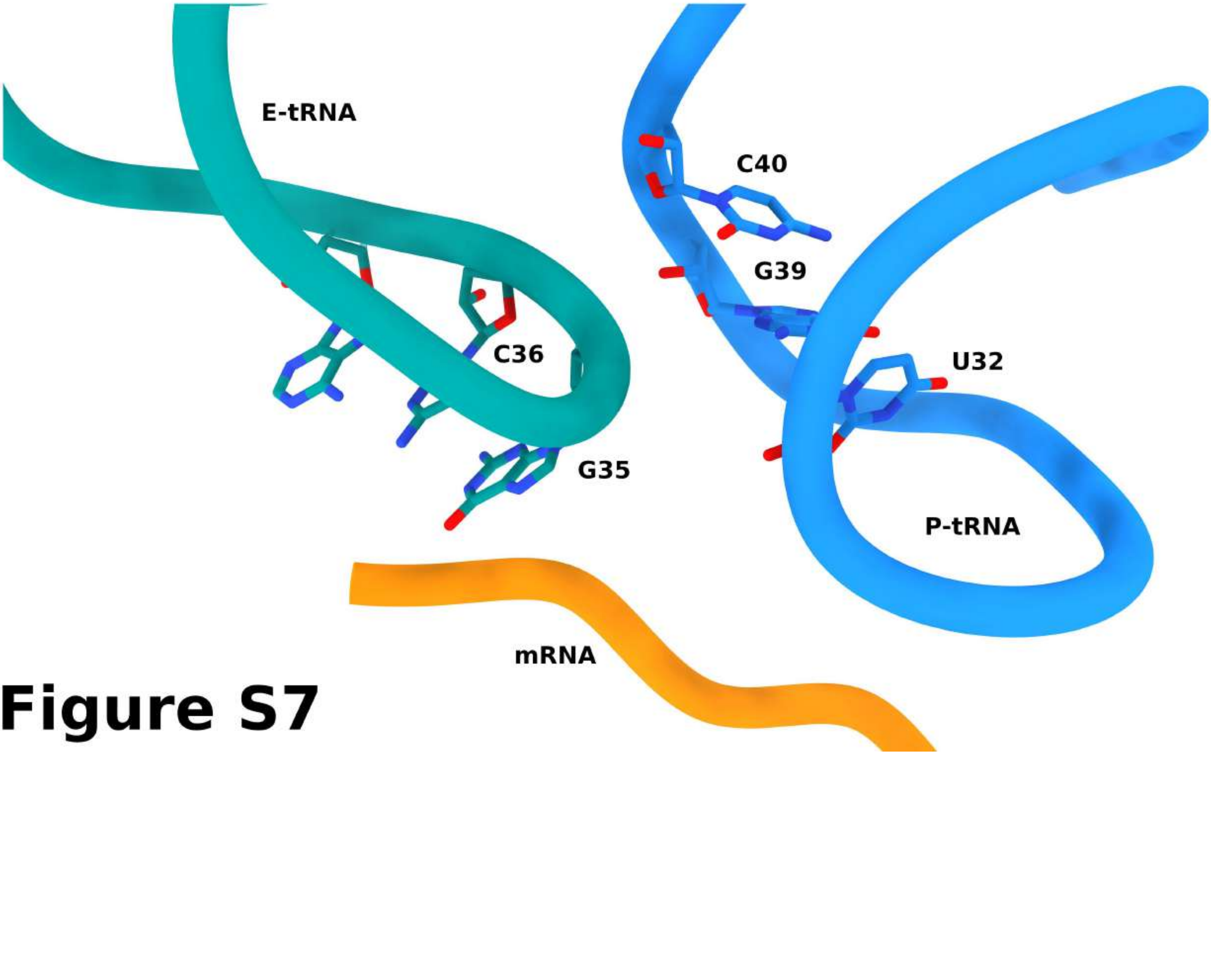


### Figure S7

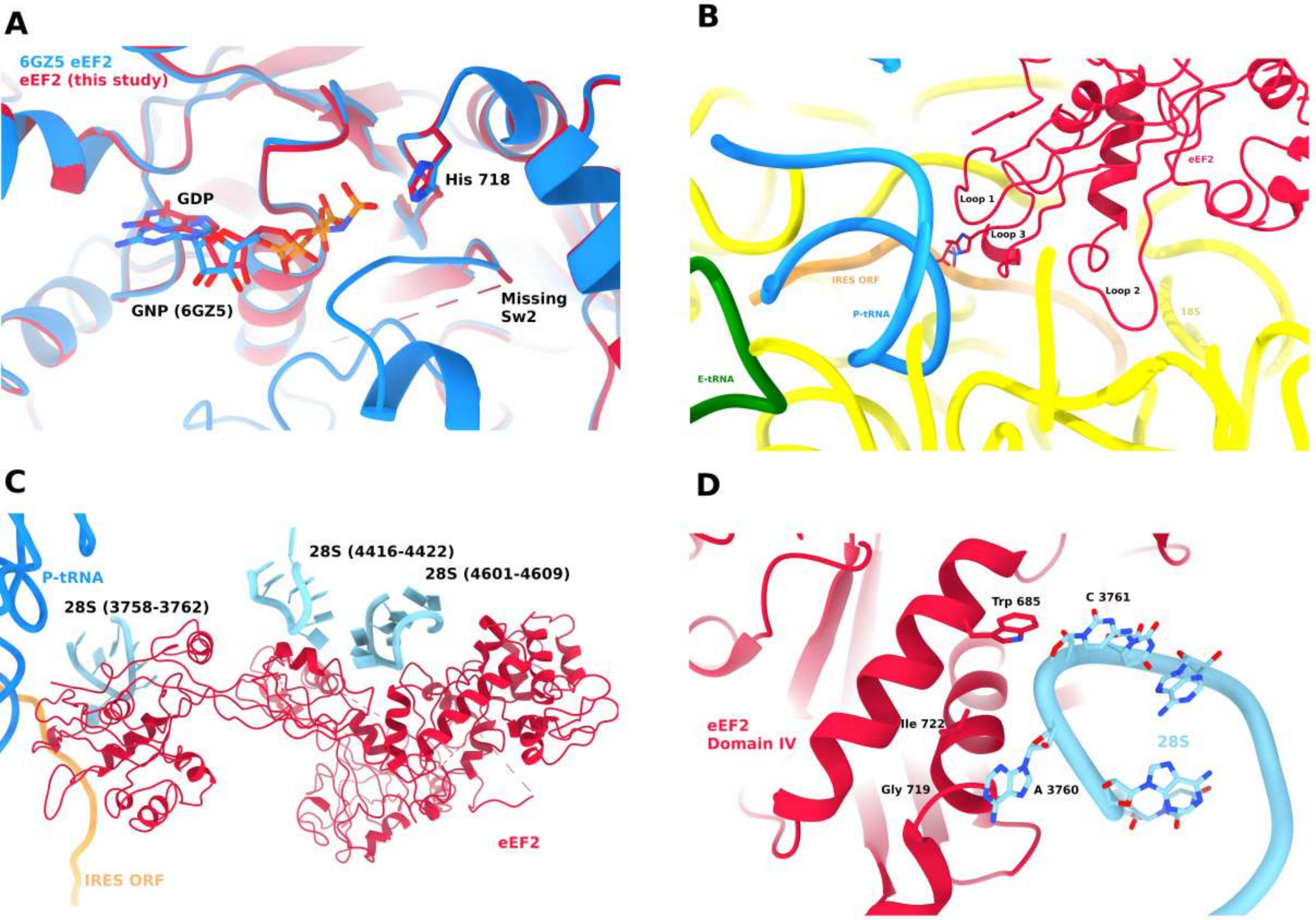
A
6GZ5 eEF2
eEF2 (this study)
GDP
His 718
GNP (6GZ5)
Missing
Sw2
B
Loop 1
Loop 3
eEF2
IRES ORF
P-tRNA
Loop 2
18S
E-tRNA
C
28S (4416-4422)
28S (4601-4609)
P-tRNA
28S (3758-3762)
eEF2
IRES ORF
D
Trp 685
C 3761
eEF2
Domain IV
Ile 722
28S
Gly 719
A 3760

### Figure S8

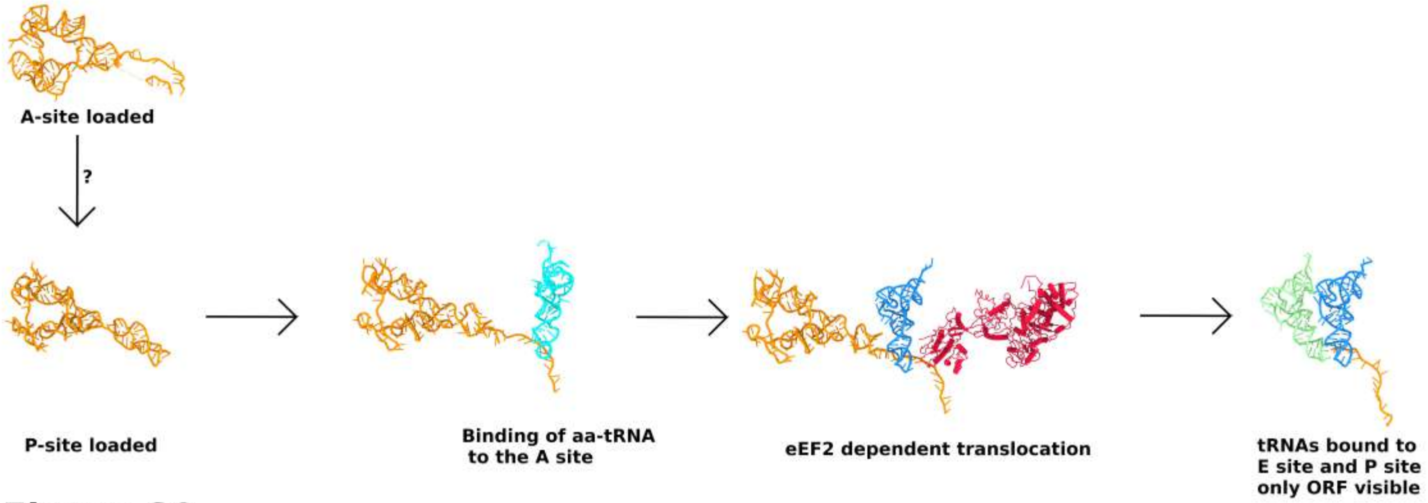
A-site loaded
?
P-site loaded
Binding of aa-tRNA
to the A site
eEF2 dependent translocation
tRNAs bound to
E site and P site
only ORF visible

**Figure S9**

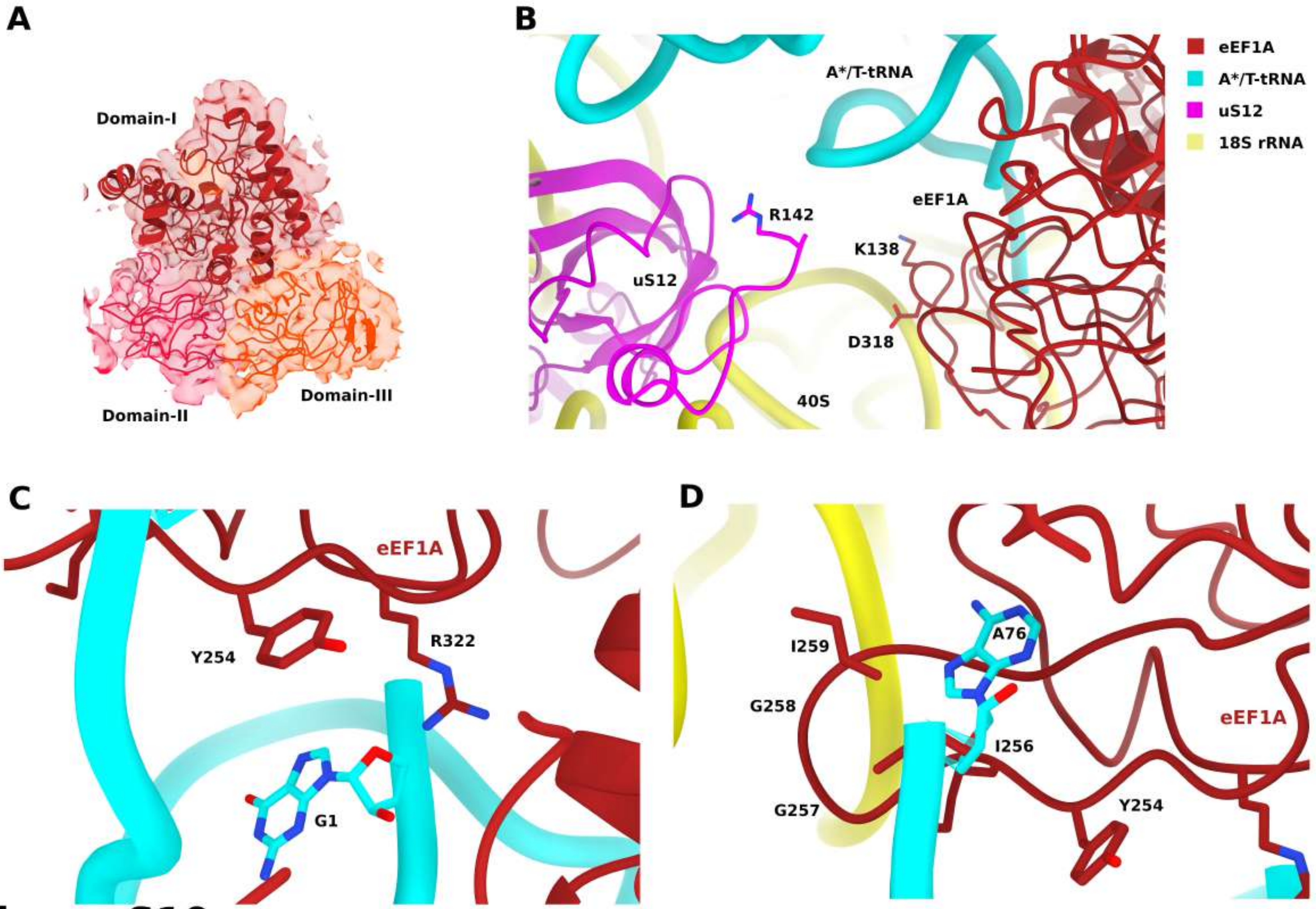
A
Domain-I
Domain-II
Domain-III
B
A*/T-tRNA
R142
eEF1A
K138
uS12
D318
40S
eEF1A
A*/T-tRNA
uS12
18S rRNA
C
eEF1A
Y254
R322
G1
D
A76
I259
G258
I256
eEF1A
G257
Y254

**Figure S10**

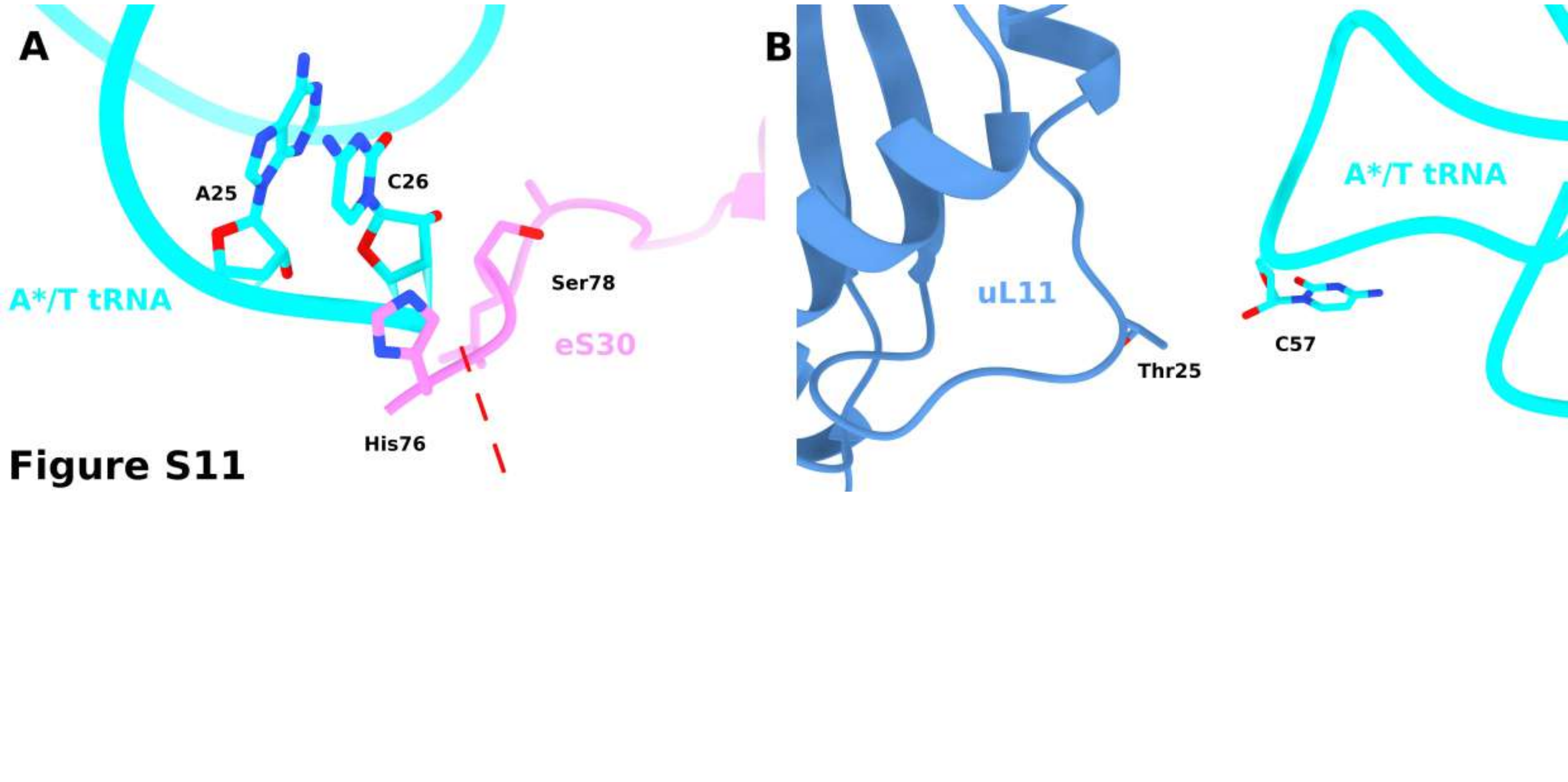


Figure S11
